# Inferring the evolving objectives that organize animal behavior

**DOI:** 10.64898/2026.09.25.754554

**Authors:** Aditi Jha, Keyue Sky Shi, Kara A. Fulton, Hanwen Lily Zhang, Christopher D. Harvey, Sandeep Robert Datta, Lisa M. Giocomo, Scott W. Linderman

## Abstract

Understanding how the brain supports flexible, goal-directed behavior is a central aim of neuroscience. Yet such behavior is highly variable and unfolds over long timescales, making it challenging to understand its organization. Advances in behavioral tracking quantify movements but do not reveal the objectives that guide them. Motivated by this, we develop inverse reinforcement learning (IRL) into a general framework for inferring these objectives from naturalistic behavior. Our approach represents an animal’s objectives as internal rewards, and infers these rewards as well as the policies they induce from observed behavior. It applies across diverse behavioral settings, from freely moving behaviors to structured tasks. It can infer objectives that switch over time or gradually change with experience. In mice capturing crickets, it identified a small set of objectives that the animals switched between over the course of a trial. In mice learning a cued labyrinth, it inferred how the animals’ internal rewards changed as learning progressed. Across both datasets, the models predicted held-out actions and generated extended behavior that reproduced key features of animal performance. By transforming long, variable behavioral sequences into time-resolved estimates of the animal’s objectives, our approach provides a quantitative basis for studying how the brain organizes natural behavior.

## 1 Main

Animal behavior is organized by the pursuit of goals in a complex and changing world. To understand the neural mechanisms that underlie this behavior, we must study animals in their natural context [1–3]. Traditionally, systems neuroscience has focused on trial-structured tasks in which goals are prescribed [4–10]. These constraints allow us to isolate specific computations and study thousands of repetitions under precise experimental control. However, they also limit the flexibility with which animals set goals, plan, and act.

The challenge with studying naturalistic behavior is that it is difficult to quantify and it rarely repeats itself. Advances in pose estimation have largely addressed the former concern, making it possible to track naturalistic behavior at scale [11–15]. The latter concern has yielded methods for behavioral segmentation and clustering, which reveal that naturalistic behavior is built from short, stereotyped movement motifs that recur across animals [16–21]. These methods reveal the structure of behavior, but not its purpose [22]. To recover that purpose, we need methods that can infer the objectives behind naturalistic behavior.

An animal’s objectives are not necessarily apparent in its movements. The same movements can serve different goals, and the same goal can be reached through many sequences of movements. Objectives may only become clear through extended behavior [23], spanning seconds to minutes, as an animal works its way toward a particular location or tracks down its prey. To infer those objectives, we must work backward from observed behavior to reason about the goals that may have motivated it [24].

Inverse reinforcement learning (IRL) addresses this problem directly [25–27]. Whereas reinforcement learning describes how rewards give rise to behavior, IRL goes in reverse, asking what reward function would make an animal’s observed behavior most likely. Given behavior and the dynamics of the environment, it recovers a reward, policy, and value function that together form an interpretable account of the objectives an animal pursues and how they organize its actions. These same variables are represented throughout the brain, where a growing body of work has related reward, value, and goal signals to circuits spanning the hippocampus, prefrontal and orbitofrontal cortex, and striatum [28–33]. Recovering them from behavior thus offers a path to connect naturalistic behavior to the neural computations that produce it.

Recent work has begun adapting IRL to animal behavior, though largely within specific task settings [34– 39]. Here we develop IRL into a general framework for studying naturalistic behavior in terms of the objectives that shape it, across tasks and over long timescales. Our formulation represents behavior at the level appropriate to each setting, from continuous trajectories in freely moving animals to discrete choices in structured tasks. It also lets the reward depend on variables that are never directly observed —the goal an animal is currently pursuing, which may switch from one moment to the next, or its knowledge of the environment, which accumulates gradually with experience.

We first validate the approach on a simulated navigation task with known rewards, then apply it to two datasets of naturalistic mouse behavior that exemplify the two cases above. In cricket capture, mice instinctively hunt over seconds to minutes. While specific movement trajectories are never repeated, the model recovers a small set of interpretable goals that persist across many movements and switch within single trials. In a labyrinth navigation task, we model each mouse from its first exposure and recover a reward that forms as the animal learns, concentrating on the rewarded locations as its knowledge of the maze sharpens. Across both datasets, our approach converts behavior into time-resolved, quantitative estimates of the objective an animal pursues, extending naturalistic behavior analysis from describing what an animal does to explaining why it does it.

## 2 Results

### Modeling behavior using inverse reinforcement learning

An animal’s objectives shape its behavior. However, whereas we can observe an animal’s movements, its objectives are not directly observable. We therefore represent these objectives as the animal’s internal rewards. We first build a generative model that describes how rewards drive behavior, and then use inverse reinforcement learning (IRL) to infer those rewards from observed behavior.

We describe behavior as a sequence of states and actions in a Markov decision process. At each timestep, the animal occupies a state *s*_*t*_, which describes its situation in the world, and executes an action *a*_*t*_, which then carries it to a new state, *s*_*t*+1_ (Fig. 1a, b).

**Fig. 1.**
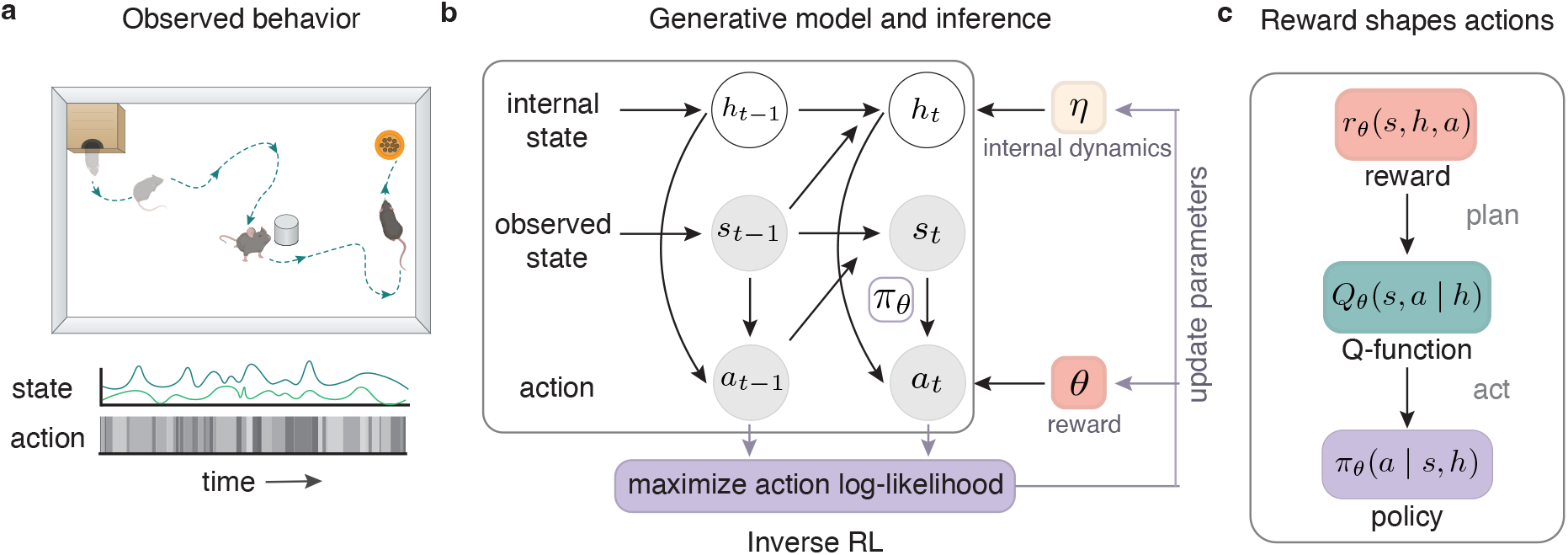
**(a) Observed behavior.** We represent behavior as a sequence of observed states *s*_*t*_ and actions *a*_*t*_. The state summarizes the animal’s situation in the world, and the action represents what the animal does next. **(b) Generative model and inference**. As the animal moves through its environment, the next state follows from its current state and action according to the environment’s dynamics. Each action depends on both the observed state *s*_*t*_ and an additional internal state *h*_*t*_. This internal state also evolves over time, depending on the previous internal state *h*_*t−*1_ and observed state *s*_*t−*1_. Inverse reinforcement learning works backward from the observed actions, and infers the reward parameters *θ* and internal-state dynamics parameters *η* that maximize the likelihood of those actions. **(c) Reward shapes actions**. The reward *r*_*θ*_(*s, h, a*) specifies the immediate value of taking action *a* in state *s*, conditional on the current internal state *h*. Planning then yields the action-value *Q*_*θ*_(*s, a* | *h*), the expected cumulative reward from taking that action. The animal then selects its action from the policy *π*_*θ*_(*a* | *s, h*), which favors actions of higher value. As *h* changes, the animal’s action preference can change even at the same observed state.

We parameterize the immediate reward for taking action *a* in state *s* as *r*_*θ*_(*s, a*), using a flexible function (a multilayer perceptron in all applications below). Planning with this reward yields the value of each action, *Q*_*θ*_(*s, a*), the expected sum of future rewards after taking action *a* in state *s* (Fig. 1c). Since animal behavior is variable, we model action selection with a softmax policy that favors higher-value actions:

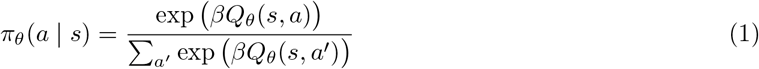

Here *β* is an inverse-temperature parameter; larger *β* concentrates probability on higher-value actions.

IRL uses this generative model to infer rewards from observed behavior [27, 40]. Each candidate reward assigns a probability to every action through the policy. We infer the reward parameters *θ* by maximizing the likelihood of the observed actions (Fig. 1b):

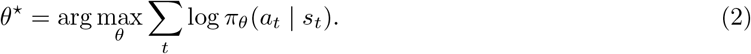

The inferred reward thus provides a quantitative account of the objectives that best explain the observed behavior under this model.

This formulation assumes that the objective remains fixed throughout a trajectory [27, 40]. For naturalistic behavior, however, this assumption is too restrictive. An animal’s action preferences may change at the same observed state depending on its internal state. We therefore condition the reward on an internal state *h*_*t*_, such as the goal the animal is currently pursuing or its accumulated knowledge of the environment, giving *r*_*θ*_(*s*_*t*_, *h*_*t*_, *a*_*t*_) (Fig. 1b, c). The animal then plans given this internal state, yielding a policy *π*_*θ*_(*a*_*t*_ | *s*_*t*_, *h*_*t*_). As *h*_*t*_ evolves, so does the reward associated with a given observed state and action, allowing the animal’s effective objective to vary over time.

Our approach is agnostic to the representation of states and actions. They can be chosen to match the behavior under study, from continuous descriptions of freely moving behavior to discrete states and choices in structured tasks. The internal state *h*_*t*_ can likewise be chosen to capture the source of variation relevant to the behavior. The principle of maximizing the likelihood of observed actions (Eq. (2)) remains the same across these choices, while state-action representations determine how the likelihood is computed and optimized (see Methods for details). We exploit this flexibility in the settings that follow.

### 2.1 IRL recovers changing goals in a simulated navigation task

An animal’s objectives and rewards are not directly observed, so we cannot test inferred objectives against ground truth from behavior alone. We therefore began with a simulation in which the true rewards were known. We designed the task with features that make naturalistic behavior hard to understand: the same objective could be achieved through multiple routes, and the objective being pursued itself changed over time.

We simulated an agent navigating a circular arena containing three curved blockades, a home site and a food site, shown in Fig. 2a. Each trial began at a random location, and at any moment the agent pursued one of three objectives, which we refer to as goals: seeking home, seeking food, or exploring the perimeter. Each goal had its own baseline transition probability. For the two site-directed goals, seeking home and seeking food, contact with the corresponding site increased the probability of leaving that goal (Fig. S1). Each goal persisted for a variable number of actions, and the active goal changed several times within a trial.

**Fig. 2.**
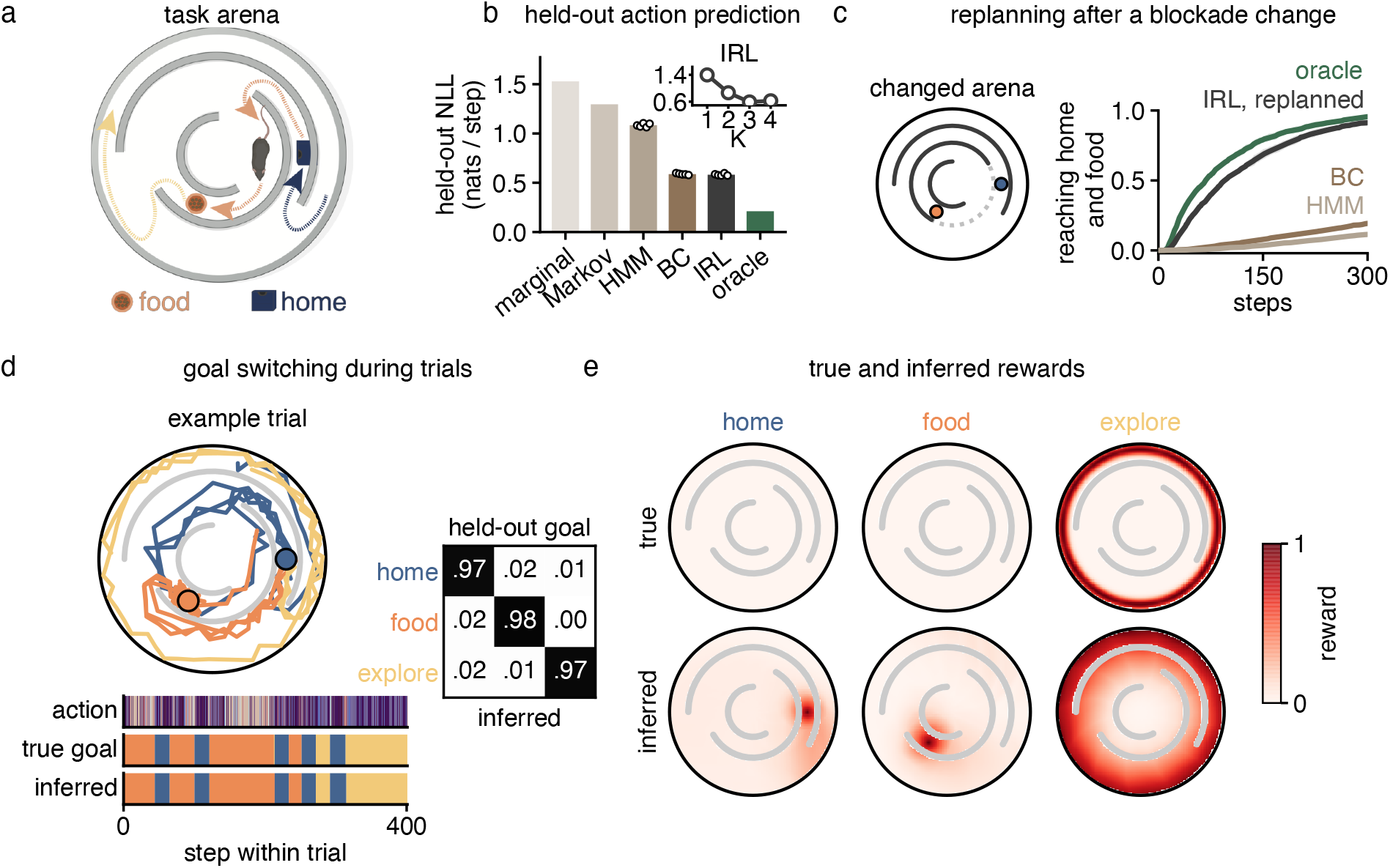
Simulated navigation task. (a) The arena, containing three curved blockades, a home site (blue) and a food site (orange). Dotted arrows illustrate paths associated with the three goals, seeking home (blue), seeking food (orange) and exploring (yellow). (b) Held-out negative log-likelihood of the next action for each model. The marginal baseline predicts each action from its overall frequency; the Markov model predicts each action from the preceding one; the hidden Markov model (HMM) allows those transition statistics to differ across latent behavioral states; behavior cloning (BC) maps the observed state directly to the next action without planning; the oracle is the generating policy. Each circle is a random seed. Lower values indicate better prediction. (c) Left, the arena after we rotated the middle blockade; the dotted arc marks the original position. Right, fraction of episodes reaching both home and food versus elapsed steps, for the oracle (green), IRL re-planned under the new dynamics with the inferred rewards held fixed (black), BC and HMM carried over unchanged (shades of brown). Shading, s.e.m. across seeds. (d) Left, an example held-out trial with the trajectory colored by inferred goal; filled circles, the two rewarding sites. Below, the action, the true and inferred goal sequences on a shared time axis. Right, confusion matrix between true and inferred goals on held-out trials, after aligning the arbitrary goal labels. (e) True (top) and inferred (bottom) reward for each goal, evaluated across the arena.

We represented the agent’s observed state by its location, as well as distances and bearings to the two rewarding sites and the arena wall. Its six actions corresponded to fixed-length movements in different directions, with execution noise. Its internal state captured the latent active goal, which determined the reward function it used for planning. Therefore, under different goals, the same location could favor different actions. Moreover, the blockades required extended routes, allowing the active goal to shape actions well before the agent reached a site.

We then fit IRL to trajectories generated by this agent. IRL jointly inferred the latent goal sequence, the goal-transition dynamics, and a reward function for each goal. Held-out negative log-likelihood stopped improving beyond three goal states, *K* = 3, matching the generating number of goal states (Fig. 2b).

We next asked how well this model predicted held-out actions relative to other approaches. A marginal baseline predicted each action from its overall frequency. A first-order Markov model predicted each action from the preceding one, following the model of syllable transitions used in MoSeq [18, 21]. A hidden Markov model (HMM) allowed these transition statistics to differ across latent behavioral states, following shMoSeq [41]. We also compared IRL with a behavior-cloning model (BC) that mapped the current observed state directly to the next action, without planning. We used the same baseline model classes for all subsequent datasets. The movement-sequence models predicted actions less accurately, consistent with their omission of the agent’s spatial context. BC, by contrast, matched IRL when given the same observations. Both models optimized action likelihood, but BC learned a direct mapping from states to actions, whereas IRL predicted actions through inferred rewards and planning.

IRL recovered both the goals and their rewards. Inferred goal sequences matched the generating ones on held-out trials (Fig. 2d). The food and home rewards were organized around their corresponding sites, and the exploratory reward favored movement along the wall (Fig. 2e). The inferred transition matrix and goal-duration distributions likewise matched their generating values (Fig. S1). Goal recovery remained accurate with only about 5% of the training data (3,200 time steps) and under the tested action-set misspecifications (Figs. S2 and S3). It deteriorated, however, when relevant environmental information was omitted from the state.(Fig. S3).

The advantage of recovering a reward rather than directly mapping states to actions is most apparent when the available routes change. We rotated the middle blockade while preserving the rewarding locations. Holding the inferred rewards fixed, we re-planned under the new arena dynamics (Fig. 2c). The re-planned IRL model reached both sites nearly as efficiently as the oracle. Despite matching IRL on held-out action prediction in the original arena, BC rarely reached both sites. Thus, IRL transferred by using the recovered rewards to plan new routes, whereas BC retained its learned state-to-action mapping and had no planning step to incorporate the changed dynamics.

Together, these simulations show that IRL can recover goals that switch within a trial, including a goal defined by a pattern of movement rather than a destination. By inferring reward from behavior, IRL also supports re-planning when the environment changes.

### 2.2 Switching goals organize cricket capture behavior in mice

We next applied our framework to mice capturing a cricket, a naturalistic behavior performed in an open arena (Fig. 3a). The arena contained a home box at a fixed location, where the mouse began each trial. A cricket was released at the opposite end of the arena, the home-box door was opened, and the mouse was free to hunt. The trial ended once the mouse captured the cricket and returned to the home box to consume it. Across four mice and 68 sessions (15–20 sessions per animal; median 22 trials per session), we recorded 1,618 trials comprising 8 hours of behavior with a median trial length of 14 s.

**Fig. 3.**
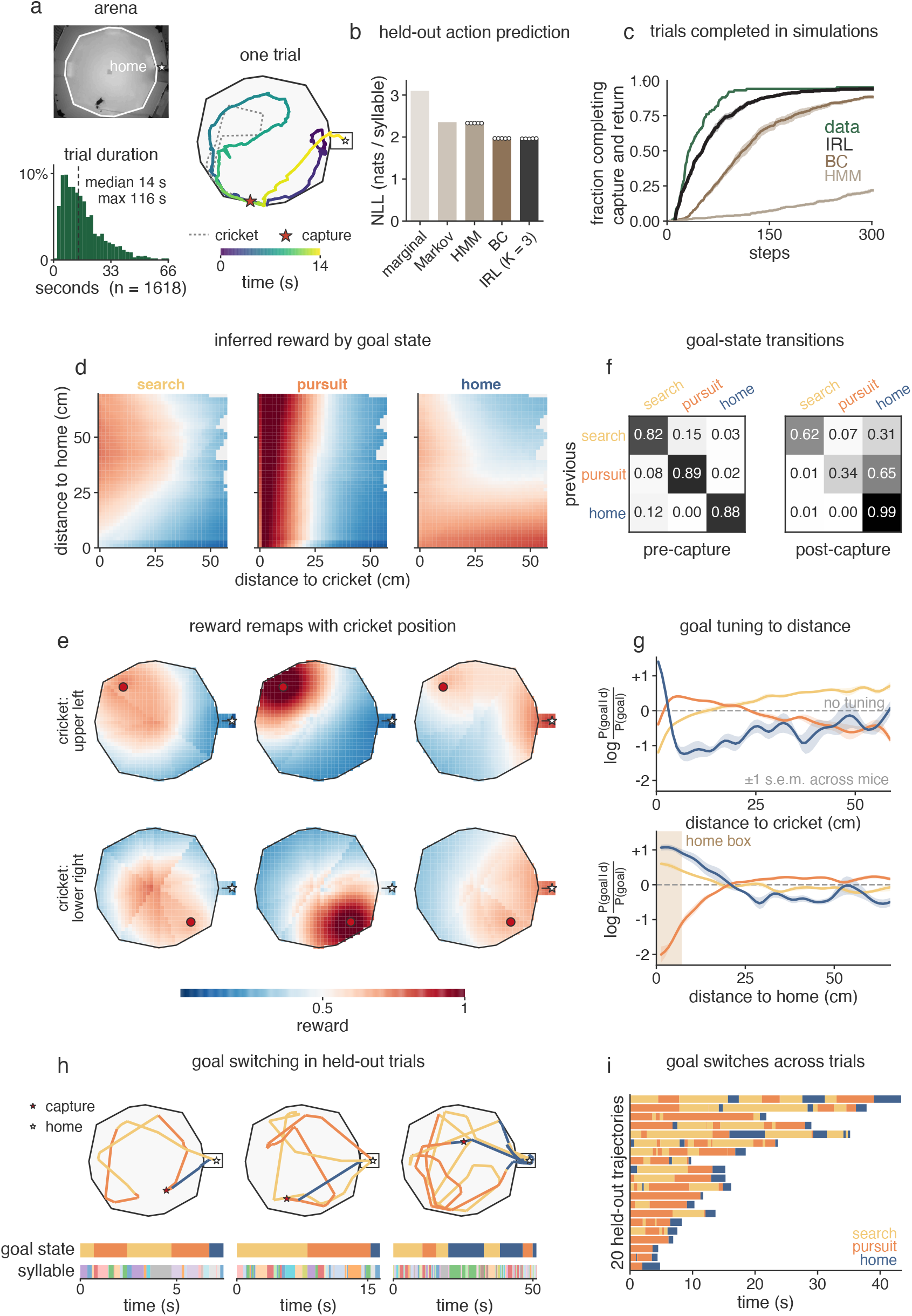
Switching goals during prey capture in mice. (a) Left, overhead frame of the arena, with the home box marked by a star, and distribution of trial durations across 1,618 trials (dashed line, median). Right, an example trial showing the mouse’s trajectory colored by elapsed time and the cricket’s trajectory (dotted line). Filled star shows when the cricket was captured; open star indicates home. (b) Held-out negative log-likelihood (NLL) of the next syllable for a marginal baseline, Markov, HMM, behavior cloning (BC), and IRL. The IRL model uses *K* = 3 goal states. Each circle is a random initialization. (c) Fraction of empirical trials and matched-condition rollouts completing both cricket capture and the subsequent return home versus elapsed steps. Green, data; black, IRL; dark brown, BC; light brown, HMM. Shading, s.e.m. (d) Inferred reward for each of the *K* = 3 goal states as a function of distance to the cricket and to the home box, with the remaining state variables marginalized. (e) The same rewards evaluated across the arena, with the cricket fixed in two quadrants (rows) and goal states in columns as in (d). Red dot represents cricket; star, home box. Color scale shared with (d). (f) Inferred goal-state transition matrices before (left) and after (right) cricket capture. Rows indicate the previous goal state and columns the next goal state. (g) Goal tuning to distance from the cricket (top) and home box (bottom), expressed as the log ratio of the posterior goal probability at a given distance to its overall probability; dashed line, no tuning (zero). Lines, mean; shading, *±* 1 s.e.m. across mice; tan region, extent of the home box. (h) Three example held-out trials: trajectory colored by inferred goal state (filled star, capture; open star, home), with goal-state and syllable sequences below on a shared time axis. (i) Inferred goal-state sequences for 20 held-out trajectories, ordered by duration. See also Videos S1–S5.

Trials varied by nearly two orders of magnitude in duration (Fig. 3a). Some trials consisted of direct runs to the cricket and back home, whereas in others the mouse lost track of the cricket, disengaged, or returned home before capture. Individual movement trajectories rarely repeated across trials. This variability poses a challenge for methods that look for structure in movement transitions alone. The next movement depends on the mouse’s changing relationship to the cricket and home, not on the preceding movement alone. We therefore asked whether IRL could explain these diverse trajectories in terms of the objectives the mice pursued.

To this end, we tracked pose from overhead video using SLEAP [15] and segmented continuous movement into 30 behavioral syllables using keypoint-MoSeq [21]. We treated each syllable as an action and represented the animal’s observed state by its distances and egocentric bearings to the cricket, home box, and arena wall. We also included the preceding syllable to capture short-timescale dependencies between successive movements.

We modeled the animal’s objective as a discrete latent goal state that could switch over time and modulate its reward. Since cricket capture changes the task from hunting to returning home, we allowed goal-transition rates to differ before and after capture. We jointly inferred the goal states, their capture-dependent transition dynamics, and the reward functions underlying each goal from behavior (see Methods for fitting details).

#### Allowing goals to switch improves prediction and generation of behavior

We first compared IRL with a marginal baseline, a Markov model of actions, an HMM over action transitions [41], and behavior cloning. The *K* = 3 IRL model achieved substantially lower held-out negative log-likelihood than the marginal and movement-sequence models and performed comparably to BC (Fig. 3b). Although the HMM also included latent states, increasing *K* did not produce a comparable improvement (Fig. S4), indicating that the gain came not from latent structure alone but from using the latent states to specify goal-dependent rewards (see Fig. S7 for comparison of inferred latent states by both methods). Within IRL, most of the improvement over a single goal was achieved with *K* = 2, with a smaller gain at *K* = 3 and little change thereafter. We therefore focus on the *K* = 3 model below; complete model-selection results are shown in Fig. S6.

Accurate one-step predictions do not necessarily produce coherent behavior over longer timescales. We simulated IRL, BC, and HMM forward from matched initial conditions and measured the fraction of rollouts that completed the trial, defined as capturing the cricket and subsequently reaching home. IRL most closely reproduced the empirical trial-completion curve, although its rollouts remained slower than the animals’ behavior (Fig. 3c). BC eventually completed most trials but did so more slowly, whereas HMM completed only about one quarter of rollouts within the simulation horizon. Thus, despite performing similarly to BC in predicting the next syllable, IRL better reproduced how often and how quickly the full capture-and-return sequence was completed.

#### Inferred rewards track task-relevant targets

Next, we examined the rewards inferred by our model. We visualized each reward as a function of distance to the cricket and home, marginalizing over the other state variables (Fig. 3d). The three rewards exhibited distinct spatial organizations. One was highest near the cricket across a broad range of distances from home (Fig. 3d, middle). A second was elevated near the home box and when the mouse was close to the cricket, consistent with the post-capture phase in which the mouse carries the cricket home (Fig. 3d, right). The third was more diffuse, with only a weak concentration around the cricket (Fig. 3d, left). We therefore refer to these states as “pursuit”, “home”, and “search”, respectively.

To determine whether these rewards tracked task-relevant targets rather than particular arena locations, we evaluated each reward at every position in the arena with the cricket fixed in two different quadrants (Fig. 3e). The pursuit reward shifted with the cricket, whereas the home reward remained anchored to the fixed home box, and the search reward remained spatially diffuse. Thus, the inferred objectives were organized around behaviorally relevant targets.

We next asked where each inferred goal was expressed during the animals’ behavior. At every held-out time point, the model provided a posterior probability over the three goals. We quantified each goal’s enrichment or depletion at different distances relative to its overall probability, averaging these profiles across mice (Fig. 3g; Methods). Pursuit was enriched at short but nonzero cricket distances and depleted farther away. Home was enriched when the mouse was within the home box, and at the smallest cricket distances, consistent with periods after the cricket had been captured. Search was depleted near the cricket and enriched farther away, with additional enrichment near the home box. Together, these spatial relationships supported the interpretation of the inferred states as search, pursuit and home.

#### Goals organize behavior over longer timescales than movements

Inferred goals persisted far longer than individual movements. Before capture, each goal had a high probability of persisting (Fig. 3f, left), with each goal lasting for 6–9 syllables. Switches occurred primarily between search and pursuit, and from home to search. The latter were consistent with the animal resuming its search after returning home without capturing the cricket. After capture, transitions from search and pursuit into home became more likely, and home almost always persisted once entered, reflecting the change in task structure at capture (Fig. 3f, right).

This separation of timescales was also apparent in individual trajectories (Fig. 3h). A single goal often persisted while the animal moved through several different syllables, yet each goal drew on much of the syllable repertoire rather than a distinct subset (Fig. S7). The inferred goals thus organized sequences composed of many different movements.

Across 20 randomly sampled held-out trajectories, goal sequences varied in both duration and composition (Fig. 3i). Search and pursuit often alternated within a trajectory, whereas home was usually concentrated near the end of the trial. Together, these results show that inferred goals persist across individual movements while switching over longer timescales during cricket capture.

Overall, IRL recovered three interpretable reward functions organized around searching, pursuing the cricket, and returning home, together with time-resolved estimates of when the mouse pursued each objective. Thus, IRL separated the objectives conserved across trials from the variable timing of their expression.

### 2.3 IRL tracks an objective as it forms during maze-learning

We next applied our framework to mice learning a cue-based navigation task. Seven animals navigated a 127-node binary-tree maze (based on [42]) with four water ports (Fig. 4a) over ten 45-minute sessions. On each trial, one of four cues was active, marked by a momentary sound and a persistent light at the corresponding port. The animal received a water reward upon arrival at that port. After an inter-trial interval of approximately 10 s, a new cue was drawn. For each animal, we concatenated the ten sessions in chronological order so that its experience accumulated across sessions.

**Fig. 4.**
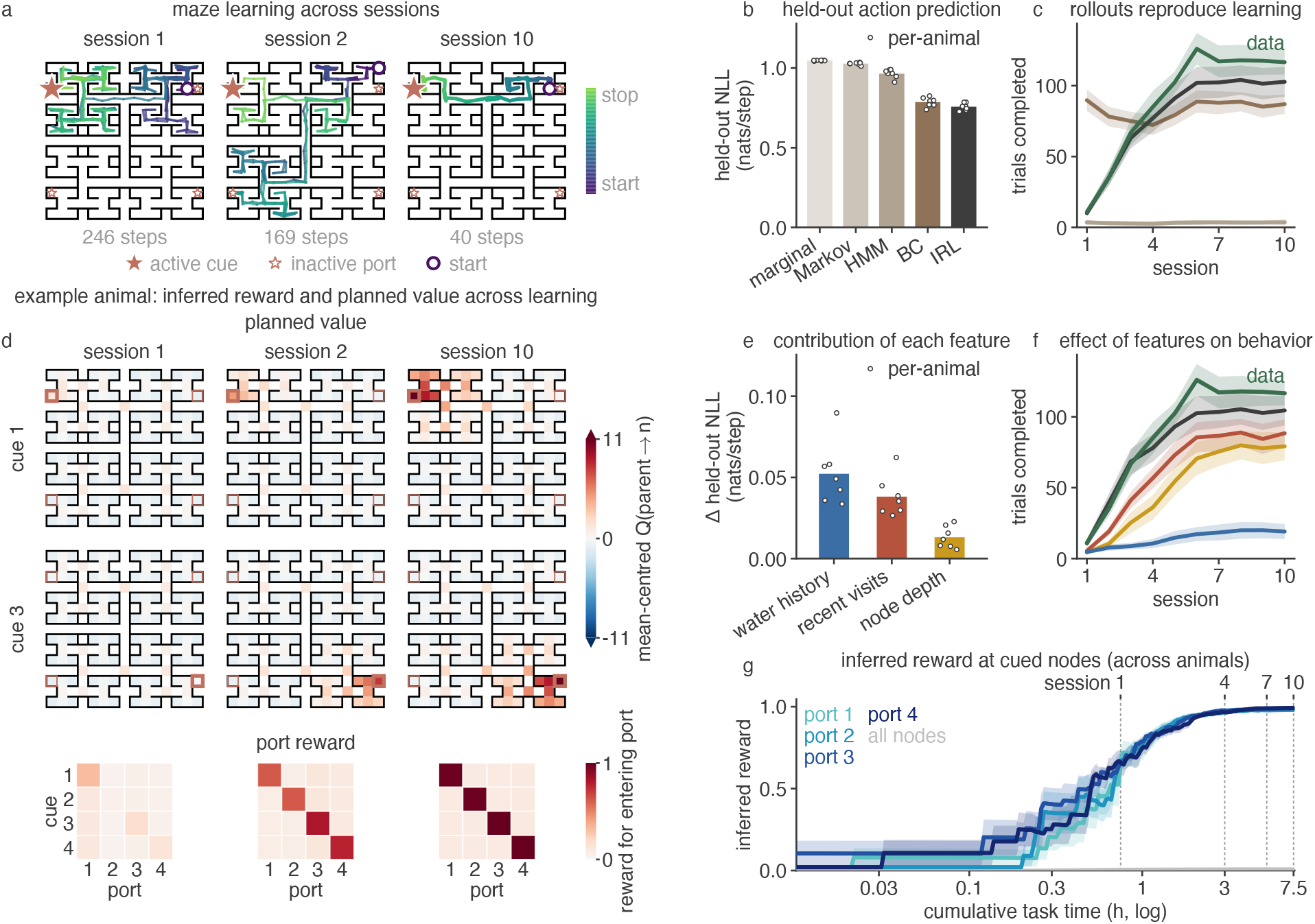
Experience-dependent IRL predicts maze learning and infers cue-specific rewards. (a) Example trajectories from one animal in sessions 1, 2, and 10, colored by elapsed time; stars, reward ports; enlarged star, port indicated by the active cue. Path length is shown below each maze. (b) Held-out negative log-likelihood of the next action for each model. Circles, individual animals; bars, mean. (c) Trials completed per session by the animals (green) and in rollouts from the models in (b). Shading, s.e.m. across animals. (d) Top, mean-centered value of entering a node for two cues (rows) in sessions 1, 2, and 10 (columns). Bottom, inferred reward for entering each port, organized by active cue (rows) and port (columns), for the same sessions. (e) Increase in held-out negative log-likelihood relative to the full model, after removing water history, recent visitation, or node depth and refitting the model. Circles, individual animals; bars, mean. (f) Trials completed per session in rollouts after the corresponding feature ablations (colors as in e); green, animal data. Shading, s.e.m. across animals. (g) Mean inferred reward for entering each port when its associated cue was active, and mean reward at other maze nodes (gray) versus cumulative task time. The time axis is logarithmically scaled; dotted lines mark selected session boundaries. Shading, s.e.m. across animals. See also Video S6.

We modeled each animal from its first exposure to the task. The task contingencies were fixed from the outset, but the animals had to learn both the maze layout and which port was rewarded under each cue. Their behavior changed with experience, from long, circuitous paths early in training to efficient navigation toward the correct port (Fig. 4a). This improvement established that mice learned the task, but did not reveal what the animals learned and how that knowledge guided their choices. We asked whether IRL could reveal how the animal’s internal rewards developed with experience.

At each step *t*, the observed state *s*_*t*_ specified the animal’s current node, the active cue, and whether it was in a trial or in an inter-trial interval. From this state, the animal selected an action *a*_*t*_ from the moves available at its current node. The resulting task outcome *y*_*t*_ indicated whether the animal obtained water. We distinguish this observed water collection from the internal reward inferred by IRL.

To model learning, we allowed the animal’s rewards and knowledge of the maze to change with experience. A reward function of only the current node and cue would assign fixed rewards throughout training, whereas fitting separate rewards for each session would not explain how they changed with experience. We therefore used a single reward function with parameters shared across sessions, but whose inputs depended on the animal’s experience.

This required a different internal state from cricket capture, within the same framework. In cricket capture, *h*_*t*_ was a discrete latent goal that switched from moment to moment. Here, *h*_*t*_ accumulated gradually and summarized what the animal had experienced before each step. It contained a visitation history tracking how recently and frequently each node had been visited, a water history tracking water received at each node under each cue, and a transition history recording the maze transitions the animal had experienced. The visitation and water histories decayed at distinct rates. Once these rates were selected, *h*_*t*_ was fully determined by the observed trajectory.

The internal reward for entering a node depended on its visitation history and its water history under the active cue, together with its depth in the maze and whether the animal was in a trial or an inter-trial interval. Each node therefore had its own reward, but computed from these features rather than from the node’s identity, so any port-specific reward had to arise from experience alone (see Methods for details).

The transition history served a different role: it determined the animal’s estimate of where each action would lead, allowing knowledge of the maze layout to develop with experience. Planning over the maze’s nodes combined these estimates with the current internal rewards to determine the action policy. Thus, visitation and water histories shaped what the animal valued, while transition history shaped how it planned through the maze.

#### Experience-dependent IRL predicts and generates maze learning

We first asked whether the experience-dependent IRL model predicted held-out actions better than the baseline models. We compared it with the action-sequence baselines: marginal, Markov and HMM, as well as a tabular BC model that learned a separate action distribution for each location and cue but did not change with experience. IRL achieved the lowest held-out negative log-likelihood, modestly outperforming BC and more substantially outperforming the action-sequence models (Fig. 4b).

We next asked whether the fitted models could reproduce the animals’ improvement over training, rather than merely predict the next action. We ran each model forward through the maze and measured the number of trials completed in each session (Fig. 4c; simulations for individual animals are shown in Fig. S8). During these rollouts, IRL updated its histories from its own simulated visits, transitions, and water outcomes rather than from the animals’ trajectories. The IRL rollouts reproduced the animals’ learning curve, closely tracking the rise in completed trials over the first sessions and plateauing slightly below the animals’ final performance. In contrast, the BC model performed relatively well from the outset but did not improve, showing that stationary location- and cue-specific action probabilities could capture aspects of behavior but not its experience-dependent change. The action-sequence models rarely completed the task. Thus, among the models tested, only experience-dependent IRL reproduced both the animals’ performance and its improvement over training.

#### The inferred objective becomes cue-specific with experience

We next examined how the inferred reward and the values produced by planning changed over training. For an example animal, we evaluated both quantities at the end of sessions 1, 2, and 10 (Fig. 4d).

The lower panels of Fig. 4d show the internal reward assigned to entering each port under each cue. Early in training, these rewards showed little cue-specific structure. By session 10, the reward under each cue was highest at its corresponding port. The upper panels show the value of entering a node across the maze. Early values were weakly organized, whereas later values were higher along routes leading toward the appropriate cue-specific ports. The reward captured which destinations the animal valued, while the values combined these rewards with its learned knowledge of the maze to identify routes toward them.

Thus, cue specificity emerged from the animal’s changing visitation and water histories. Evaluating the inferred reward throughout the trajectory provided a continuous readout of this process (Fig. 4g). Across animals, the reward assigned to the cue-associated ports increased relative to the reward at other nodes, with most of the separation developing during the first two sessions. The model thus provided a time-resolved account of how the objective guiding each animal’s behavior formed through experience.

#### Water history and spatial features support learning

We next asked which components of the model supported its performance. We removed water history, visitation history, or node depth information and refitted each model. Each ablation increased held-out negative log-likelihood (Fig. 4e), with cue-specific water history contributing most.

The same ablations had distinct effects on the behavior the model generated (Fig. 4f). Without cue-specific water history, performance remained near zero and did not improve across sessions. Removing recent visitation or node depth preserved improvement over training but slowed learning and lowered final performance. Learning therefore depended on the history of cue-specific water outcomes, while recent visitation and maze depth shaped how efficiently the animal navigated. Accounting for transition history during planning improved action prediction primarily in the early sessions, compared with assuming that the animals knew the maze layout from the outset (Fig. S9).

Overall, experience-dependent IRL inferred an internal reward that became organized around the cue-specific ports, produced values that guided navigation through the maze, and reproduced learning in rollouts using only the model’s own accumulated experience. Together with the cricket-capture results, this shows that the framework can track both objectives that switch within ongoing behavior and objectives that develop gradually through learning.

## 3 Discussion

We developed an inverse reinforcement learning framework to infer the objectives that organize naturalistic behavior. In natural contexts, animal behavior is shaped by self-directed objectives that are not directly observable, and that may change as the animal acts [43, 44]. We represent them with a reward function, and infer this function along with the policy it induces. This yields an interpretable generative model of behavior. The model’s scope depends on three representations. The observed state describes the animal and its surroundings. Actions define the unit of choice, such as a behavioral syllable during a hunt or a turn at a maze junction. The internal state captures a latent goal or accumulated experience. It also shapes the reward function, allowing the same observed situation to lead to different choices as goals or experience change. By tailoring these representations to the behavior under study, our approach applies across distinct settings.

In simulations, IRL recovered the generating rewards and their changes over time. When the structure of the environment was changed, the inferred rewards supported efficient re-planning under the changed dynamics without refitting. This transfer shows that the recovered rewards captured objectives that extended beyond the routes used for fitting [45].

In cricket capture, inferred goals organized successive movements into periods of search, pursuit, and returning home. IRL predicted held-out actions more accurately than movement-sequence models. While behavior cloning matched its one-step prediction accuracy, rollouts generated from the model completed the task more slowly. Thus, similar accuracy in predicting individual actions did not guarantee similar behavior over long timescales [46, 47]. In the maze, inferred rewards became increasingly concentrated at cue-associated ports as experience accumulated, accounting for the animals’ shift from circuitous exploration to efficient navigation. The fitted IRL model also reproduced the animals’ learning curve using its own simulated experience.

Previous work has used IRL to study visuomotor behavior [48, 49], thermotaxis in *C. elegans* [35], and changing goals in tasks with discrete state spaces [36–39]. Related inverse control methods have been demonstrated with task-specific reward or cost parameterizations [50–52]. Building on these, our framework combines flexible reward functions with evolving internal states to model behavior across settings.

By converting long, variable behavioral sequences into time-resolved estimates of goals, rewards, and values, this framework provides a quantitative account of the objectives that organize behavior and candidate variables for relating those objectives to neural activity [31, 53, 54]. Inferred goal transitions provide alignment points in self-directed behavior, in the absence of an external cue that marks when the animal’s objective changes [41]. These transitions let us ask how neural activity changes when goals switch [30], and whether activity associated with a goal persists across the different movements used to pursue it. Tracking inferred rewards and values during learning similarly lets us ask how neural activity changes as an objective takes shape [55]. Including movement, observed state, and these inferred variables in the same analysis would test whether inferred objectives explain additional variation in neural activity [56, 57]. This offers a route to studying how the brain organizes naturalistic behavior, even when animals do not repeat the same movement sequence.

## Limitations

First, our inferences depend on the state, action, and internal-state representations we choose. These representations are modeling assumptions that determine which sources of behavioral variation the model can express. If we omit a relevant variable, the model may fail to account for its effects or absorb its influence into the inferred reward or internal state.

Second, we rely on a model of environmental dynamics to infer rewards. Errors in those dynamics, or a mismatch between them and the animal’s own model of the environment, can distort the rewards we recover [58].

Third, we cannot generally identify a unique reward function from behavior alone, since different rewards can produce the same policy [27, 59, 60]. Likewise, the interpretability of an inferred latent state does not establish that it corresponds uniquely to the animal’s subjective goal. We therefore interpret the inferred rewards and internal states as behaviorally grounded explanations under our modeling assumptions, rather than uniquely determined accounts of what the animal values.

Fourth, we assume that the animal plans conditional on its current internal state, without anticipating how that state will change. Allowing such anticipation would require including the internal state in the planning dynamics, and when the animal is uncertain about relevant variables, planning over beliefs about those variables [61].

## Acknowledgments

We thank Camila Blank for her work on initial preprocessing of the maze datasets. We thank Caleb Weinreb, Benjamin Cowley and members of the Linderman Lab for helpful discussions. We also thank Hannah Lee, Henry Smith and Yoel Sanchez Araujo for feedback on the manuscript. AJ was supported by the Wu Tsai Interdisciplinary Postdoctoral Fellowship. KAF was supported by the Yang Tan Collective at Harvard, Hock E. Tan and K. Lisa Yang Center for Autism Research at Harvard. SRD was supported by the Simons Collaboration on the Global Brain, the Simons Collaboration on Plasticity and the Aging Brain, and NIH grants 1RF1NS135504, 1R01NS136355, and 1R01DC021965. LMG is an HHMI Investigator. This work was supported by the National Institutes of Health 1R01MH126904-01A1 (LMG), R01MH130452 (LMG), BRAIN Initiative U19NS118284 (LMG), The Vallee Foundation (LMG), The James S. McDonnell Foundation (LMG), The Simons Foundation 542987SPI (LMG), and The Howard Hughes Medical Institute (LMG). SWL is supported by grants from the NIH (U01NS136507, R01NS131987, RF1MH133778, R01NS130789, & R01AG097491), the NSF (2440859), and the Sloan, Simons, and McKnight Foundations.

## Methods

### Experimental model and study participant details

#### Mouse subjects (cricket capture)

Experimental subjects were 2 female C57BL/6J and 2 male C57BL/6J mice aged 3-5 months, obtained from Jackson Laboratory. All animal experiments were conducted in accordance with the guidelines and regulations of the National Institutes of Health and Stanford University under APLAC protocol 27694. Mice were housed on a reverse 12-h light cycle at 19–23^*°*^C and 30–70% humidity, and experiments were performed during the dark phase. Following Implant surgery, mice were singly housed with a running wheel. After 3d post-op recovery, mice were food restricted to 85-90% of baseline weight throughout the duration of behavioral assay to ensure sufficient motivation for eating crickets during recordings.

#### Mouse subjects (maze)

Maze navigation experiments were approved and carried out in accordance with Harvard Medical School Institutional Animal Care and Use Committee (IACUC). Experimental subjects include five male and two female C57BL6/J mice harboring the Vglut2-flp and Tbx21-cre alleles (Jackson Laboratory stock no. 030212; B6;129S-Slc17a6tm1.1(flpo)Hze/J and Jackson Laboratory stock no. 024507; B6;CBA-Tg(Tbx21-cre)1Dlc/J). All mice were recorded in the maze between 5-7 months of age and maintained on a 12h:12h reverse light cycle with food ad libitum. Mice were habituated to handling and recorded during the dark cycle under soft white light and infrared illumination. Mice were water-restricted 5 days prior to behavioral testing and throughout the duration of the behavioral assay to ensure sufficient motivation for navigating the maze.

### Method details

#### Inverse reinforcement learning to infer rewards from behavior

To model animal behavior, we describe each behavioral episode as a trajectory *ζ* = (*s*_1_, *a*_1_, …, *s*_*T*_, *a*_*T*_), where *s*_*t*_ *∈ S* denotes the animal’s observed state, *a*_*t*_ *∈ A*(*s*_*t*_) its action, and *T* the trajectory length. Its behavior unfolds through the interaction with the environment, whose transition dynamics are given by *T*(*s*^*′*^| *s, a*) = *P* (*s*_*t*+1_ = *s*^*′*^| *s*_*t*_ = *s, a*_*t*_ = *a*). We assume that the animal’s action selection is shaped by an unknown underlying reward function *r*_*θ*_(*s, a*) *∈* ℝ with parameters *θ ∈* ℝ ^*D*^. Inverse reinforcement learning (IRL) then seeks to infer the reward parameters *θ* that make the observed behavior most likely. Below, we describe our reward inference procedure. Throughout, we take the action space *A* to be discrete and finite.

#### Maximum-entropy inverse reinforcement learning

We use maximum-entropy IRL [27, 40], which assigns greater probability to trajectories that accumulate greater reward while otherwise maximizing the entropy of the trajectory distribution:

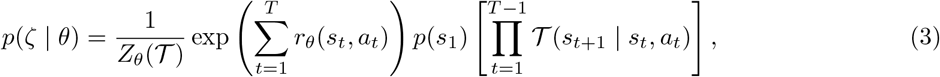

where *Z*_*θ*_ is the normalizer obtained by summing over all feasible trajectories. This formulation natually accommodates variability in animal behavior, as it spreads probability across trajectories rather than committing to a single optimal one.

An equivalent policy representation of Eq. 3 makes explicit how the reward shapes each action [62]. We use a discounted, temperature-scaled form of this policy. A discount factor *γ ∈* [0, 1) sets the effective planning horizon, weighting future rewards as 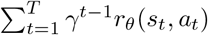 [63, 64]. An inverse temperature parameter *β >* 0 controls how deterministically the animal acts on its value [65]. The resulting policy is:

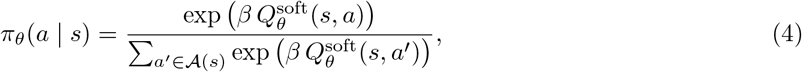

where 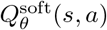 is the entropy-regularized expected cumulative reward starting from (*s, a*). It satisfies the following soft-Bellman equations:

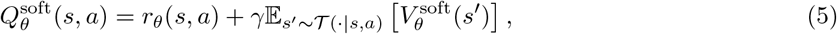

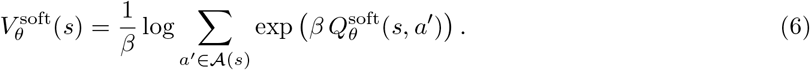

Eq. 4 makes clear that the reward specifies a distribution over available actions at each state. This allows us to infer *θ* by maximizing the likelihood of the observed actions under the induced policy,

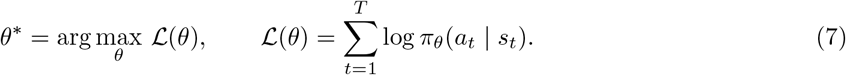

Fitting alternates between computing the policy, and updating the reward parameters to increase the overall likelihood of actions. Both steps depend on the state representation. For small discrete state spaces, we use dynamic programming and differentiate through its updates. For continuous states, we approximate the policy with a neural network and estimate the reward updates from simulated trajectories. We describe both in detail below.

#### Discrete state spaces

When is *S* finite and small enough to enumerate, we solve the soft-Bellman equations exactly by dynamic programming, sweeping Eqs. 5–6 to convergence over all states and actions. Every operation in this recursion is differentiable, so we obtain *∇* _*θ*_*ℒ* by backpropagating and optimize Eq. 7 directly by stochastic gradient ascent.

#### Continuous state spaces

When *S* is continuous, we cannot enumerate the states or solve the soft-Bellman equations exactly. Instead, we alternate between computing the policy induced by the current reward and updating the reward using simulated trajectories [66, 67].

For a fixed reward *r*_*θ*_, we approximate the soft action-value function with a neural network 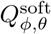. We fit its parameters *ϕ* using soft *Q*-learning [65]:

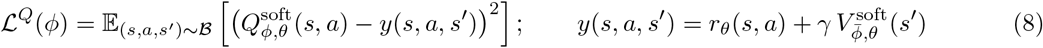

Here 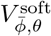 is computed using Eq. 6, with a slowly updated copy 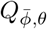 of the *Q*-network. The network induces an approximate policy *π*_*ϕ,θ*_ through Eq. 4. At each step of the optimization, we simulate this policy in the task environment, store the resulting transitions (*s, a, s*^*′*^) in a replay buffer, and optimize Eq. 8 using minibatches ℬ drawn from this buffer. The reward parameters *θ* remain fixed during this fit.

We next update the reward using the likelihood gradient. For fixed transition dynamics *T*, the gradient of the soft action value can be expressed as an expected sum of discounted reward gradients:

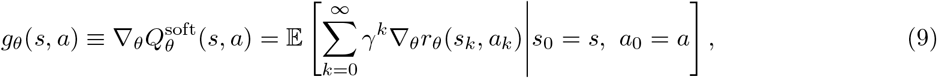

where trajectories begin with (*s, a*) and subsequently follow the transition dynamics *T* and policy *π*_*θ*_. Differentiating the log-policy then gives

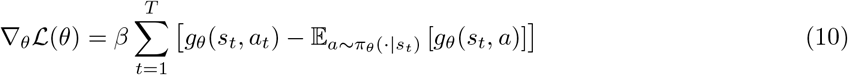

This gradient contrasts the accumulated reward gradients following the observed action with their average over actions drawn from the current policy at the same state. Thus, updating *θ* in this direction adjusts the reward to make the observed actions more likely.

We estimate these quantities using the fitted policy. For each observed state *s*_*t*_ and available action *a*, we generate *L* simulated rollouts that start with *a* and subsequently follow *π*_*ϕ,θ*_. Each rollout terminates after *H* steps or upon reaching a terminal state. Averaging the discounted reward gradients across these rollouts gives *ĝ* _*θ*_(*s*_*t*_, *a*). Since our action space is discrete, we compute the policy average in Eq. 10 by summing over all actions 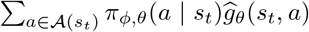. Substituting this average and *ĝ* _*θ*_(*s*_*t*_, *a*_*t*_) into Eq. 10 gives the sampled likelihood gradient 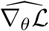.

Fitting therefore alternates between policy fitting (*ϕ*) and reward updates (*θ*). The reward updates use the simulated reward gradients, and thus do not differentiate through the soft-*Q* fitting procedure.

#### Rewards modulated by internal state

An animal’s objectives may switch over time or change with experience. To capture this, we introduce an internal variable *h*_*t*_ *∈ ℋ*, and let the reward, *r*_*θ*_(*s*_*t*_, *h*_*t*_, *a*_*t*_), depend on it. Parameters 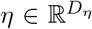 govern how *h*_*t*_ evolves over time. Depending on the behavior, *h*_*t*_ may be a discrete latent goal that switches over time [37–39], or a deterministic summary of the animal’s experience. We use the former in simulated navigation and cricket capture and the latter in the maze task.

In our setting, *h*_*t*_ modulates the reward without expanding the space over which the animal plans. In other words, the animal plans under its current internal state without anticipating how that state will change. At each time *t*, we hold *h*_*t*_ fixed, and plan over the observable state space *S*,

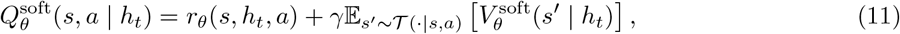

yielding *π*_*θ*_(*a* | *s, h*_*t*_).

Inference over *h*_*t*_ depends on how it is parameterized. When *h*_*t*_ is constructed deterministically from the animal’s experience, it can be computed from the observed history for a given *η*. We then maximize the conditional action log-likelihood.

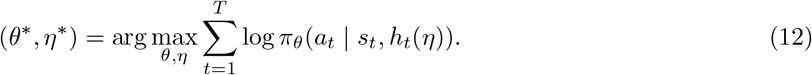

When *h*_*t*_ is instead a latent goal *z*_*t*_ *∈ {*1, …, *K}*, we model it as a state-dependent Markov chain,

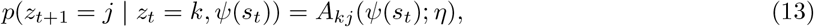

where *ψ*(*s*_*t*_) contains observed state features that influence goal switching, and *η* parameterizes the transition model. In our applications, *ψ* encodes a small number of task conditions, reducing Eq. 13 to a separate transition matrix for each condition. More generally, *ψ* may contain other observed features. We denote the initial goal distribution by *p*_0_ *∈* Δ^*K−*1^, with *p*_0_(*k*) = *p*(*z*_1_ = *k*).

We then maximize the trajectory likelihood marginalized over possible goal sequences:

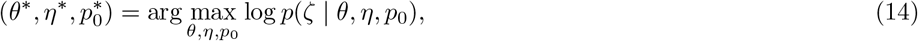

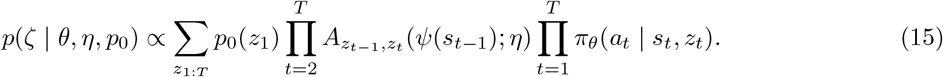

The omitted initial observed-state *p*(*s*_1_) and environment-transition factors *p*(*s*_*t*+1_ | *s*_*t*_, *a*_*t*_) do not depend on *θ, η*, or *p*_0_. For multiple independent trajectories, we maximize the sum of their log-likelihoods.

We optimize Eq. 15 by generalized expectation-maximization. Given the current parameter estimates, (*θ, η, p*_0_), log *π*_*θ*_(*a*_*t*_ | *s*_*t*_, *k*) provides the emission log-likelihood for goal *k*. The forward-backward algorithm then returns

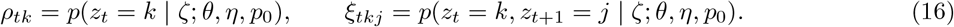

For *N* independent trajectories, we update the initial goal distribution as 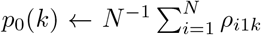, where *ρ*_*i*1*k*_ is the posterior probability that trajectory *i* begins in goal *k*. We update *η* using the posterior transition probabilities and the corresponding state features. Each goal’s reward is updated using the action log-likelihood weighted by its posterior responsibilities. Algorithms 1 and 2 summarize our fitting procedure.

#### Reward parameterization and identifiability

In some of our applications, rewards depend only on the state reached by an action. These rewards can still be expressed in the state–action form used above. Writing the arrival-state reward as 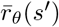 and suppressing its dependence on *h*_*t*_, we define

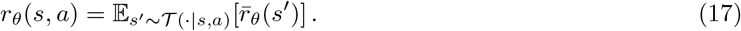

Next, reward functions are not uniquely determined by behavior. In particular, their overall scale is not separately identifiable from *β*. In all our applications, we restrict rewards to [0, 1] and let *β* set their effective scale.

We next describe the three applications: simulated navigation, cricket capture, and cued-maze navigation. We then describe the baseline models and evaluation metrics.

##### Algorithm 1

Generalized EM for IRL with latent goal states

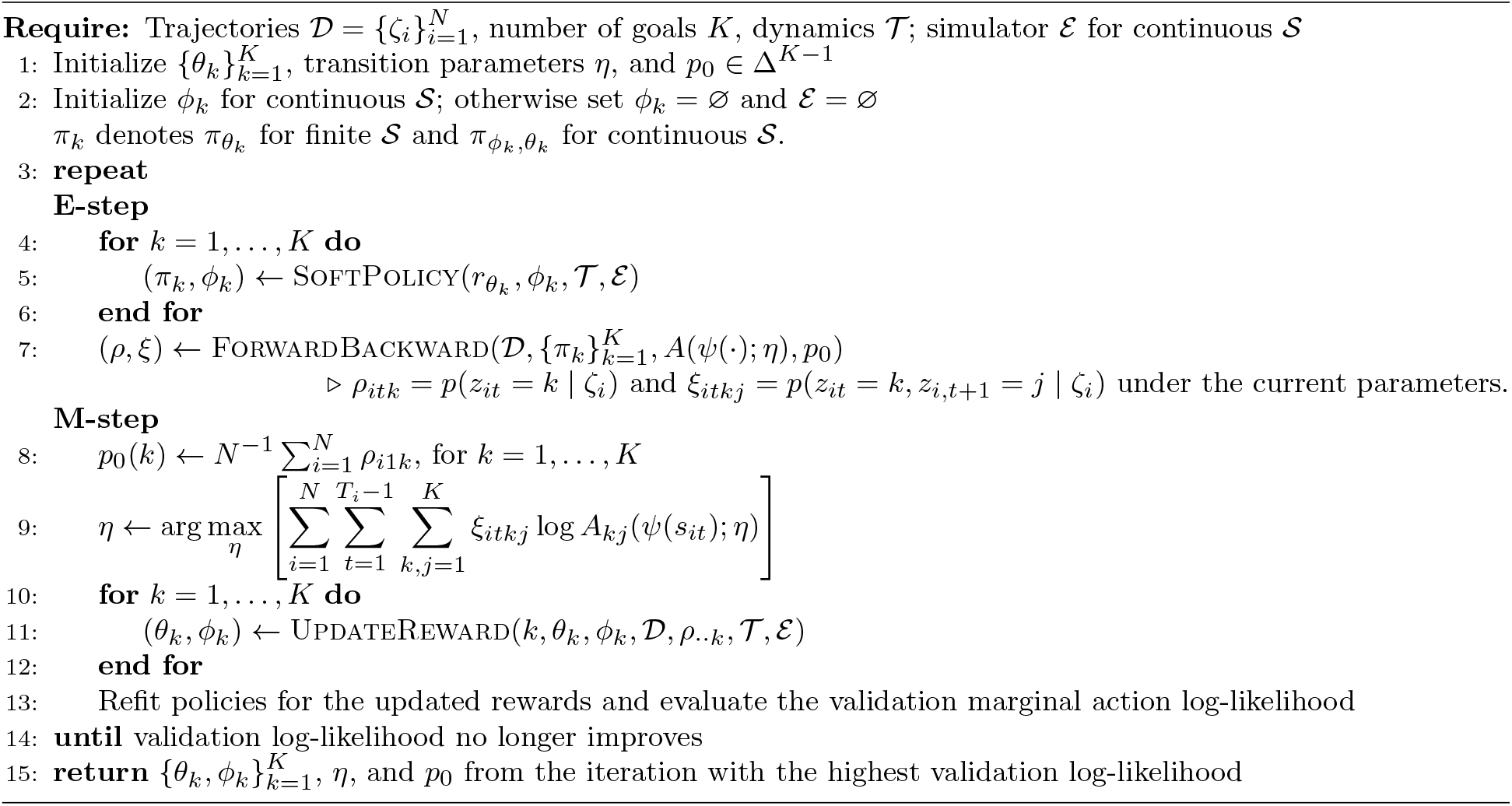

##### Algorithm 2

Reward update for one goal (UpdateReward)

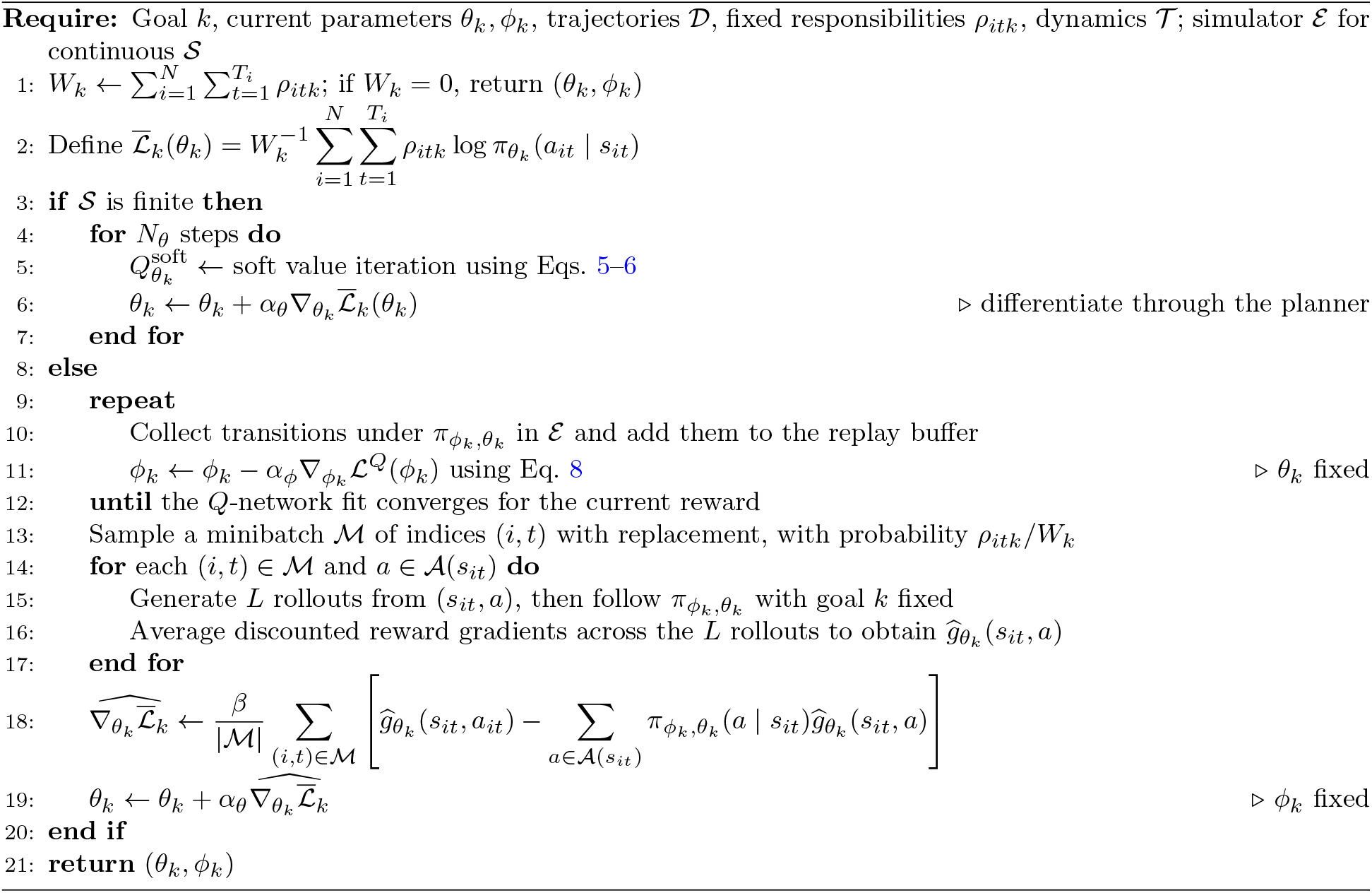

### Simulated navigation task

#### Simulated environment

We simulated a circular arena of radius 40 cm, placing a home site at (26, 0) cm, a food site at (*−* 8.5, *−* 14.7) cm, and three impassable arcs at radii 12, 22 and 31 cm (Fig. 2a). We gave each arc a span of 210^*°*^ and offset it 120^*°*^ from the next, so that reaching either site required extended detours. We let the agent’s location vary continuously in R^2^ and allowed six egocentric movements at 60^*°*^ intervals, each a step of 5.2 cm with Gaussian execution noise of s.d. 4^*°*^ in heading and 0.25 cm in length. A direction was available only if the noiseless step advanced the agent at least half a step.

#### Generating agent

The generating agent had three possible goals, seeking home, seeking food, or exploring, each modulating its reward. Under the site-directed goals we rewarded it with 1 whenever a step landed within 2.6 cm of the corresponding site (home or food) and zero otherwise. Under exploration we rewarded thigmotaxis, movement along the arena wall,

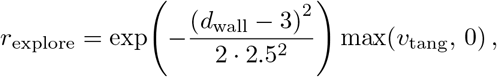

where *d*_wall_ is the distance in cm from the step’s arrival point to the arena wall, and *v*_tang_ *∈* [*−* 1, 1] is the cosine of the angle between the step and the counterclockwise tangent to the wall at that point.

The latent goal *z*_*t*_ evolved according to a state-dependent Markov chain (Eq. 13), where *ψ*(*s*_*t*_) indicated no contact, contact with home or contact with food. With goals ordered as (home, food, explore), in the absence of site contact, the per-step probabilities of leaving each of the three goals were 1*/*400, 1*/*400, and 1*/*56, respectively. When the agent contacted the site it was pursuing, home or food, the probabilities of the three goals at the next time step became (0.775, 0.1125, 0.1125) and (0.05, 0.925, 0.025), respectively. Thus, goals could switch spontaneously, but reaching an active home or food target increased the probability of leaving that goal. Complete transition matrices are shown in Fig. S1.

We obtained one policy per goal by fitting a separate soft *Q*-network to its known reward using soft *Q*-learning (Eq. 8), with discount factor *γ* = 0.95 and inverse temperature *β* = 10. Given the active goal, the agent selected actions from the corresponding policy. We refer to this three-goal generating process as the oracle.

#### Data generation

We began each episode from a position sampled uniformly over the arena, outside the two site discs, and sampled its initial goal uniformly. We generated 200 episodes of 400 steps each, holding out 25% for model selection and evaluation.

#### Model fitting and selection

We represented the observed state using 11 features: normalized *x* and *y* position of the agent, distances to the home site, food site, and arena wall, and the sine and cosine of its egocentric bearing to each. Positions were normalized by the radius of the arena. The action specified one of the six egocentric movement directions.

We treated the goal as a discrete latent state *z*_*t*_ *∈ {*1, …, *K}*. The transition features *ψ*(*s*_*t*_) indicated whether the agent contacted home, contacted food, or contacted neither site, reducing Eq. 13 to three transition matrices. For each condition *c* and current goal *k*, we updated the transition probabilities as 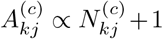. Here, 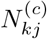 denotes the expected number of transitions from goal *k* to goal *j* under condition *c*. We normalized the transition probabilities 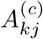 over *j*.

We parameterized each goal’s reward as a separate multilayer perceptron *r*_*θ k*_ (*s*^*′*^) evaluated on the arrival state (Eq. 17). We fitted the model using the IRL procedure in Algorithms 1 and 2, with continuous observed states *s* and the true environment as the simulator ℰ.

We evaluated *K ∈ {*1, 2, 3, 4} across five random initializations. The mean held-out action log-likelihood across the five fits for each *K* was highest at *K* = 3 (Fig. 2b, inset). We therefore used the best *K* = 3 fit for subsequent analyses. Remaining hyperparameters are given in Table 1.

**Table 1.** Simulated navigation task: model and fitting parameters. Braces denote grids swept during selection, with the selected value in parentheses.

|  |  |
| --- | --- |
| *Data* |  |
| episodes, steps each | 200, 400 |
| held out | 25% of episodes |
| *Representation and architecture* |  |
| $\dim s_t, \vert \mathcal{A}\vert $ | 11, 6 |
| $K$ | $\{1, \dots, 4\}$ (3) |
| $r_{\theta_k}$ | MLP $11 \rightarrow 32 \rightarrow 1$, one per goal |
| $Q_{\phi_k, \theta_k}^{\text{soft}}$ | MLP $11 \rightarrow 128 \rightarrow 128 \rightarrow 6$, one per goal |
| $\eta$ | $\mathbb{R}^{3 \times K \times K}$, plus $p_0 \in \Delta^{K-1}$ |
| *Planning and optimization* |  |
| $\gamma$ | $\{0.7, 0.8, 0.9, 0.95\}$ (0.95) |
| $\beta$ | $\{1, 2, 10, 20, 100\}$ (100) |
| $H$ (rollout horizon) | 32 |
| $L$ (rollouts per state-action pair) | 40 |
| $\vert \mathcal{M}\vert $ (reward-update minibatch) | 2,048 |
| $\vert \mathcal{B}\vert $ ($Q$-learning minibatch) | 512 |
| $\alpha_\theta, \alpha_\phi$ | $3 \times 10^{-3}, 1 \times 10^{-3}$ |
| replay buffer, target update | $4 \times 10^5$, Polyak 0.005 |
| seeds | 5 per setting |

#### Recovery of goals and rewards

Goal labels are arbitrary up to permutation, so we first matched inferred to generating goals by the permutation maximizing the correlation between their rewards on held-out states. We then assigned each held-out step to its maximum-posterior goal using the posterior from the forward-backward algorithm, 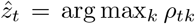, and reported the confusion matrix against the generating goal (Fig. 2d). To compare rewards directly, we evaluated the generating and inferred rewards for each goal on a grid of arena positions (grid size 0.6 cm). At each position, we retained the maximum reward across 12 arrival headings for visualization (Fig. 2e).

We also compared inferred and generating goal-transition matrices and evaluated goal recovery with a three-state HMM (Fig. S1).

To assess data requirements, we fitted models to nested subsets of 8–150 training trials, with five random seeds per subset and fixed network architectures. We evaluated all fits on the same 50 held-out trials using action NLL, reward MSE at arrival states, and recall averaged across goals (Fig. S2).

We tested model misspecification using the same 150 training and 50 held-out trials, with three seeds per condition (Fig. S3). To test state misspecification, we removed distance and bearing features for one or both sites, or for the wall, together with allocentric position. To test the influence of the transition dynamics, we varied the agent’s assumed step length from 0.25 to 2 times its true value. Finally, to test how action specification affected goal recovery, we altered the action set in several ways. We rotated the six movement directions by 20^*°*^, or used three, four, or five equally spaced directions, adjusting the planning dynamics accordingly. Reward error was the reduction in *R*^2^ relative to the correctly specified fit. For altered action sets, we matched each held-out action to the nearest model move. We split each move’s probability equally among the original actions mapped to it, then computed NLL. These are shown in Fig. S3.

#### Blockade-shift transfer

To test whether the inferred rewards transferred to new dynamics, we rotated the middle blockade by 180^*°*^ while leaving the sites and rewards unchanged. For IRL, we held the inferred rewards, goal-transition matrices, and *p*_0_ fixed and refitted only the *Q*-networks in the modified arena. We re-planned the generating model in the same way to provide an oracle reference. BC and HMM were transferred without refitting.

We generated 300 episodes of 300 steps per model from the training initial-condition distribution and measured the fraction that reached both sites. We repeated the procedure across five independent refits and reported the mean *±*1 s.e.m. across refits (Fig. 2c).

### Cricket capture

#### Behavioral task

Mice hunted crickets in a twelve-sided arena (mean center-to-vertex radius 38.2 cm, center-to-wall 36.6 cm) containing a home box on the perimeter. There were two possible home-box locations, separated by 90^*°*^ from each other. All mice underwent a single switch in home-box location partway through the experiment, and the first post-switch session was excluded. At the beginning of each trial, a cricket was placed at a random location at the far end of the arena from the home box, and the door to the home box was opened. The trial continued until the animal captured the cricket and returned to the home box, where it consumed the cricket. We restricted analysis to trials that followed this protocol. In all, we analyzed 1,618 trials from four mice, recorded in 68 sessions over 38 days (15–20 sessions per animal; median 22 trials per session, range 5–43), resulting in 8 hours of behavior. Trials lasted a median of 14.3 s (range 1.8 to 116.3 s).

#### Tracking and behavioral preprocessing

We tracked the animal’s pose from overhead video acquired at 60 Hz using SLEAP [15]. We tracked 11 body keypoints: the nose, neck, two points along the spine, tail base, left and right sides of the head-stage holder, forelimbs, and hips. We also tracked the cricket’s centroid, body start and body end. We downsampled the resulting keypoint trajectories to 30 Hz and segmented them into discrete behavioral syllables using keypoint-MoSeq [21]. We used a seven-dimensional latent space, sufficient to explain 90% of the variance in the keypoint trajectories, and allowed the model to infer up to 30 syllable states. All 30 syllables were occupied, and no post hoc merging was applied.

We evaluated *κ ∈ {*10^4^, 10^5^, 10^6^*}*, where *κ* controlled the persistence of syllables, and chose *κ* = 10^5^ to match the timescale of the behaviors we sought to capture. For each value of *κ*, we performed ten fits with random initializations and selected the model with the highest expected marginal likelihood. In the selected model, syllables lasted 0.40 s on average, with mean durations across syllables ranging from 0.13 to 1.03 s.

We treated each syllable execution as one step of the decision process. The executed syllable defined the action *a*_*t*_ at step *t*, and we evaluated the animal’s state *s*_*t*_ at syllable onset. Each trial was therefore represented as a trajectory (*s*_1_, *a*_1_, …, *s*_*T*_, *a*_*T*_) from the opening of the home box through capture and return home. Syllable duration was not modeled explicitly.

#### States, actions and environment

We represented the animal’s observed state by its egocentric relationship to the cricket, home box, and arena wall, together with its preceding syllable. Let *d*_*x*_ and *α*_*x*_ denote the distance and bearing from the animal to *x ∈ {c, h, w}*. Here, *c, h* and *w* represent the cricket, the home box, and the nearest arena wall. We defined

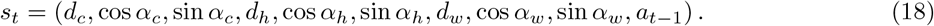

The action *a*_*t*_ identified which of the 30 syllables the animal executed. We included the preceding syllable *a*_*t−*1_ to capture short-timescale dependencies between successive movements that the spatial variables alone do not explain. Within each session, we translated coordinates so that the arena-boundary centroid lay at the origin, rotated them to place the active home box on the positive *x*-axis, and scaled them so that the home box lay at (1, 0).

Fitting the model required a simulator of the environment ℰ, which we assembled in advance and held fixed during IRL fitting. It comprised the arena geometry and home-box location for each session, together with the transition dynamics of the environment *T* (*s*^*′*^| *s, a*). The transition dynamics determined how the current state and action changed the mouse’s and the cricket’s state at the next time step.

Let *m*_*t*_ denote the animal’s position and heading and *x*_*t*_ the cricket’s position. Conditional on the executed syllable, we factorized the physical transition dynamics of the environment as

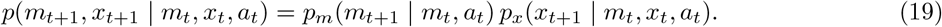

We estimated both factors from the observed behavioral sequences.

For the animal-movement factor *p*_*m*_, we measured the animal’s egocentric displacement and change in heading from the onset of each syllable to the onset of the next and fit *p*(Δ*x* | *a*) = *N* (Δ*x*; *µ*_*a*_, Σ_*a*_) and 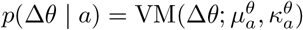 across all executions of syllable *a*.

For the cricket-motion factor *p*_*x*_, we trained an MLP to predict the cricket’s next position from the current positions of the animal and cricket and the animal’s current syllable (two hidden ReLU layers with 64 units each; two-dimensional output). We validated the cricket-motion model on held-out trial segments before capture, measuring one-step position error. We also generated 10-step rollouts by feeding predicted cricket positions back into the model while supplying recorded mouse states and syllables (Fig. S5).

The simulator combined these factors to advance the animal and cricket. It set the syllable-history component of *s*_*t*+1_ deterministically to *a*_*t*_. We additionally estimated the empirical syllable-transition probabilities 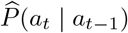 and treated syllables with zero probability as unavailable.

#### Goals and rewards

We represented the animal’s internal state as a discrete latent goal *z*_*t*_ *∈ {*1, …, *K}* with initial distribution *p*_0_. Goal transitions depended on the observed state through the distance to the cricket:

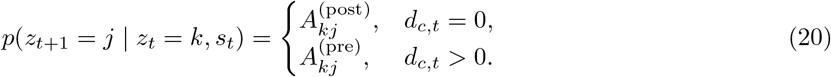

After capture, we set the distance to the cricket, *d*_*c,t*_, to zero for the remainder of the trial. Thus, we learned separate transition matrices for the post-capture and pre-capture portions of each trial, respectively. Within each regime, goal transitions were otherwise independent of the observed state.

We parameterized each goal’s reward as a separate multilayer perceptron taking the observed state *s*_*t*_ and returning one value per syllable, and passed its output through a logistic function so that *r*_*k*_(*s, a*) *∈* [0, 1]. Network architecture, remaining settings, and hyperparameter choices are given in Table 2.

**Table 2.** Cricket capture: model and fitting parameters. Braces denote grids swept during selection, with the selected value in parentheses.

|  |  |
| --- | --- |
| *Data* |  |
| trials | 1,618 |
| held out | 20% of trials |
| *Representation and architecture* |  |
| $\dim s_t, \vert \mathcal{A}\vert $ | $39 = 9 + 30, 30$ |
| preceding syllable $a_{t-1}$ | 30-dimensional one-hot vector |
| $K$ | $\{1, \dots, 5\}$ (3) |
| $r_{\theta_k}$ | MLP $39 \rightarrow 64 \rightarrow 30$, one per goal |
| $Q_{\phi_k, \theta_k}^{\text{soft}}$ | MLP $39 \rightarrow 128 \rightarrow 64 \rightarrow 30$, one per goal |
| $\eta$ | $\mathbb{R}^{2 \times K \times K}$, plus $p_0 \in \Delta^{K-1}$ |
| cricket dynamics | MLP with two 64-unit ReLU hidden layers |
| *Planning and optimization* |  |
| $\gamma$ | $\{0.7, 0.8, 0.9\}$ (0.9) |
| $\beta$ | $\{1, 2, 5, 10\}$ (5) |
| $H$ (rollout horizon) | 64 |
| $L$ (rollouts per state-action pair) | 1 |
| reward-update samples | $1,024 \times 30$ |
| $\vert \mathcal{B}\vert $ ($Q$-learning minibatch) | 128 |
| $\alpha_\theta, \alpha_\phi$ | $10^{-3}, 10^{-3}$ |
| replay buffer, target update | 50,000, every 4 ($\tau = 0.05$) |
| EM iterations | 60 |
| seeds | 5 per setting |

#### Model fitting

We pooled training trials across animals and fit a single set of goal-dependent rewards, pre- and post-capture goal-transition matrices, and an initial goal distribution, with *D* = {*ζ*_1_, …, *ζ*_*N*_*}* the pooled dataset. Because the state space is continuous, we fit the model by the sample-based procedure in Algs. 1– 2, using the estimated environment above as the simulator ℰ.

We held out 20% of whole trials and used the marginal action log-likelihood on this set throughout to assess convergence, to choose among random initializations, and to select hyperparameters. We used five random initializations per setting and swept *K ∈ {*1, …, 5} along with the grids in Table 2. Held-out likelihood improved substantially from *K* = 1 to *K* = 2, increased slightly at *K* = 3, and changed little thereafter, so we used the *K* = 3 model for analyses in the main text. Model selection results and inferred rewards for *K* = 2 and *K* = 4 are shown in Fig. S6.

#### Analysis of inferred goals and rewards

The reward *r*_*θ*_(*s, a*) depended on the animal’s spatial state, the syllable it executed, and the syllable that preceded it. To visualize it as a function of the animal’s position we first reduced over the two syllable arguments. For each spatial configuration and preceding syllable, we took the highest reward available over possible next syllables, and then averaged over preceding syllables weighted by their pooled empirical frequencies. This yielded one reward per spatial configuration, using the same weighting for every goal.

We then evaluated this quantity in two ways. For the distance maps in Fig. 3d, we averaged it over all observed states at a given distance to the cricket and to the home box, which marginalized over the animal’s position and heading within each distance pair. For the arena maps (Fig. 3e) we fixed the cricket at a chosen location, placed the animal at each position in the arena, and averaged over headings. In both cases we masked configurations the animals never occupied and used a common color scale across goals.

To ask how each goal was tuned to distance, we related goal occupancy to distance using the posterior probabilities *ρ*_*itk*_ from held-out trials. For each animal *m*, we pooled its held-out trials and estimated goal occupancy using a Gaussian kernel *K*_*σ*_ with bandwidth *σ* = 2.5 cm:

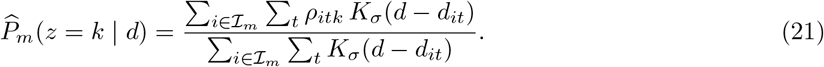

Here, *ℐ*_*m*_ denotes the held-out trials from animal *m*. We averaged these animal-level profiles across animals to obtain 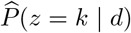. We expressed goal tuning as

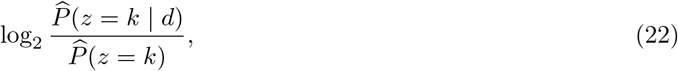

where 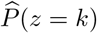 was the overall posterior goal occupancy, averaged across animals with equal weight. Positive values indicated greater goal occupancy than baseline, and negative values indicated lower occupancy. We evaluated these profiles at 64 distances evenly spaced between the 1st and 99th percentiles of the observed distances, for both distance from the cricket and distance from home (Fig. 3g).

Next, we assigned each held-out step to its maximum-posterior goal, 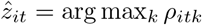. Since each step corresponded to one syllable, we expanded each goal assignment over that syllable’s duration to express the inferred sequence in time. We plotted goal and syllable sequences together for individual trials (Fig. 3h). We also plotted goal sequences for 20 randomly selected held-out trials, aligned to trial onset and ordered by duration (Fig. 3i).

For Fig. S7, we estimated syllable usage within each goal by weighting held-out syllable counts by posterior goal probabilities and normalizing within goals. We also trained multinomial logistic regressions to predict the maximum-posterior IRL goal from either the current syllable alone or the current and previous syllables. Finally, we compared IRL goals with the states of a three-state HMM. We matched HMM states to IRL goals using correlations between their posterior probabilities on training trials.

### Cued-maze navigation task

#### Behavioral task

Mice navigated the binary-tree labyrinth introduced by [42], comprising 127 nodes, with 63 binary branch points and 64 terminal nodes. We equipped four terminal nodes, one in each corner of the maze, with a water reward port, infrared beam break, and cue LED. The port locations were fixed across animals and sessions. We water restricted the mice beginning 5 days before their first maze exposure and throughout training. We recorded behavior beginning with each animal’s first exposure to the maze and followed learning over 10 training sessions, one per day, ending each session after 250 completed trials or 45 min, whichever came first. Behavior was self-paced and continuous. At the start of each trial we played a 10-kHz tone for 1 s while illuminating the LED above the target reward port, and delivered a 6 *µ*L droplet of water containing 1 *µ*M acesulfame potassium (Ace-K) at that port. The water remained available at the port until collected. We sampled the target port uniformly from the three ports other than the target on the preceding trial. Mice began each trial from their current location and could move freely through the maze with no fixed starting location. A trial ended when the animal entered the target port and broke the infrared beam to collect the reward. We separated trials by a 2-s post-reward timeout followed by an inter-trial interval of 10 *±* 2 s, during which the cue LEDs were off and the animal moved freely.

#### Tracking and behavioral preprocessing

We recorded behavior from below with a monochrome camera at 66 Hz (912 *×* 848 pixels) and tracked nose and tail-base positions using DeepLabCut [11] with a ResNet-50 network. We used the nose position only for subsequent analyses, assigning each video frame to the maze node containing the tracked nose. To turn these into trajectories for the IRL framework, we collapsed consecutive frames assigned to the same node, so that each entry in the resulting sequence corresponded to a transition. Each animal’s trajectory was an ordered sequence of moves (*n*_*t*_, *a*_*t*_), where the actions available at *n*_*t*_ were moves to the left child, the right child, and the parent, whichever existed. This discarded dwell duration at each node.

We corrected tracking artifacts before model fitting. In particular, we removed frames containing nose displacements greater than 100 px frame^*−*1^ or positions outside the maze (1.73% of frames). When this produced a gap between non-adjacent nodes, we reconstructed the intervening trajectory using the unique shortest path through the maze graph, allowing at most five intermediate nodes; reconstructed nodes accounted for 7.5% of transitions in the final dataset. We included all sessions from the seven analyzed animals, yielding 6,274 trials across 70 sessions and 293,334 node transitions.

#### Observed states, actions and internal states

We modeled each animal independently, concatenating its 10 sessions into a single behavioral trajectory that included both trials and inter-trial intervals and so covered the animal’s entire experience in the maze. At step *t*, the animal was at node *n*_*t*_, selected an available action *a*_*t*_ *∈ A*(*n*_*t*_), and arrived at *n*_*t*+1_ = *T* (*n*_*t*_, *a*_*t*_), where *T* denotes the transition structure of the maze. We let *c*_*t*_ denote the active cue, which remained fixed throughout a trial, *ι*_*t*_ *∈ {*0, 1*}* indicate whether step *t* occurred during an inter-trial interval (ITI), and *y*_*t*_ *∈ {*0, 1} indicate whether the animal collected water on arrival at *n*_*t*+1_. Note that *y*_*t*_ is an observed outcome of the task, distinct from the internal reward, which we define below.

We let the animal’s observed state be its position and current task context,

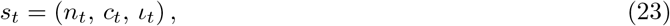

comprising its current node, the active cue, and whether it is in a trial or an ITI. Its internal state *h*_*t*_ then summarized its experience:

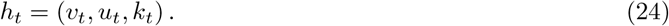

Here *v*_*t*_ *∈* ℝ^*N*^ is a visitation trace over the *N* = 127 nodes, *u*_*t*_ *∈* ℝ^4*×N*^ is a water-received trace containing one vector per cue, and *k*_*t*_(*n, a*) counts how many times the animal has taken action *a* from node *n*. The traces *v*_*t*_ and *u*_*t*_ determined the animal’s current reward, whereas *k*_*t*_ determined its current model of the maze’s transition structure.

For fixed decay rates *λ*_*v*_ and *λ*_*u*_, all three components of *h*_*t*_ were deterministic functions of the animal’s observed history. Although they represented internal quantities, they could be reconstructed exactly from the data rather than inferred as latent variables. They updated after every action as

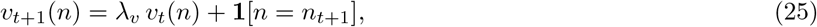

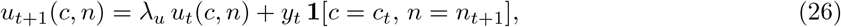

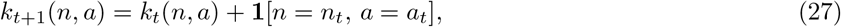

The decay rates *λ*_*v*_, *λ*_*u*_ *∈* [0, 1] were included in the internal-state dynamics parameters *η* and controlled how rapidly visitation and water histories decayed, respectively. We initialized all traces and counts to zero at the start of the first session and carried them forward across sessions within an animal.

#### Reward

At time *t*, the reward for arriving at node *n* was

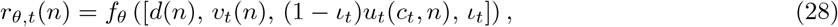

where *d*(*n*) is the node’s normalized depth in the maze and *f*_*θ*_ is a multilayer perceptron with a logistic output (Table 3). The reward therefore depended on node depth, visitation history, water received under the current cue, and ITI status. During ITIs, the water-received input was set to zero. The logistic output restricted rewards to [0, 1], with *β* setting their effective scale.

**Table 3.**
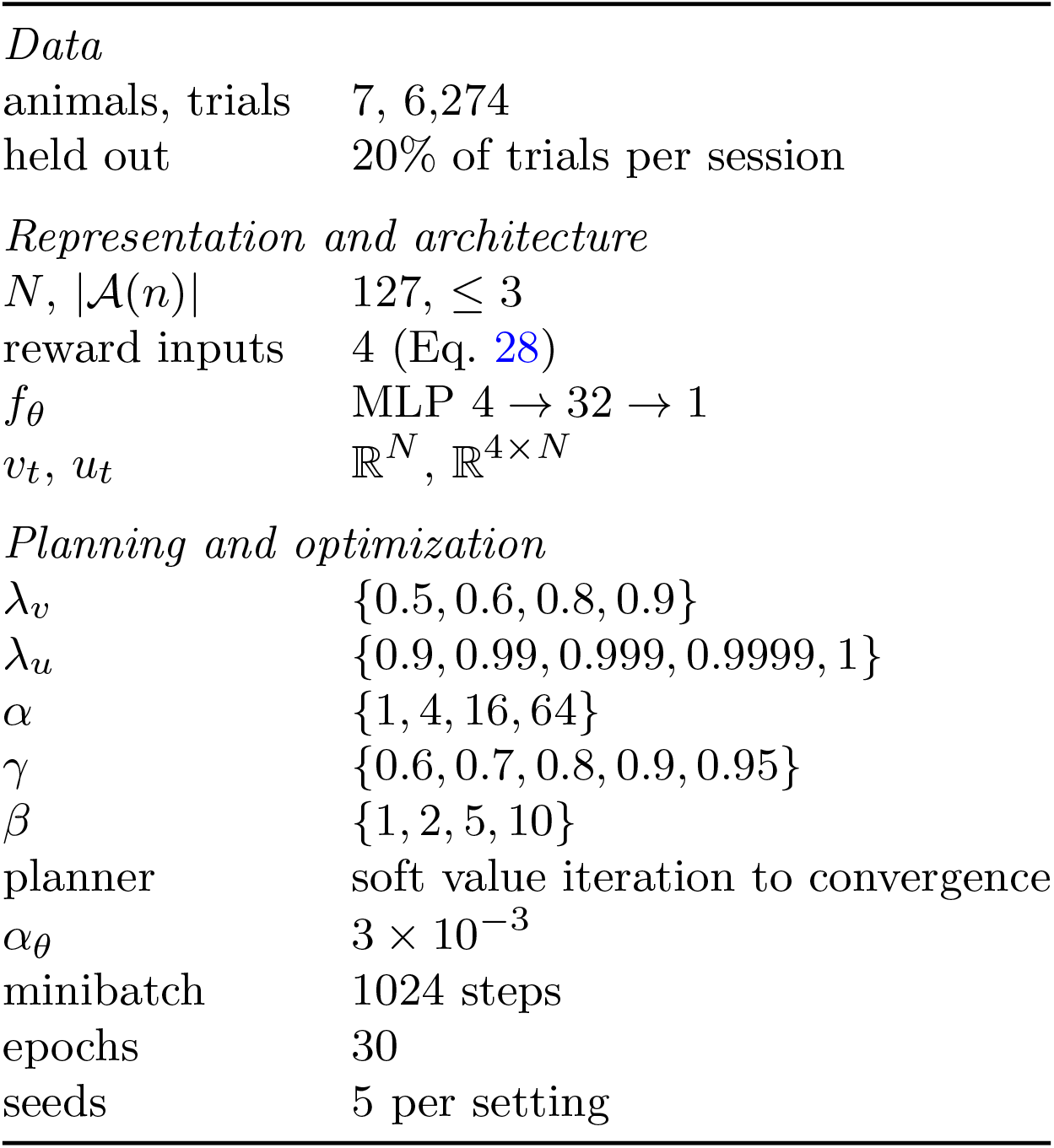
Cued-maze navigation: model and fitting parameters. Braces denote grids swept during selection.

Within each animal, the same parameters *θ* were used across all nodes and sessions. Rewards changed as the animal accumulated experience and the task context changed. We excluded node identity to prevent the reward model from encoding port locations independently of experience. Nodes at the same depth therefore differed in reward only through their experience traces.

#### The animal’s model of the maze

We did not assume the animal knew the maze’s layout, *T*, at the outset, and instead let its transition model sharpen with experience. Placing a symmetric Dirichlet prior with total concentration *α >* 0 over the successors of each node–action pair, the posterior mean transition model after *k*_*t*_(*n, a*) traversals was given as:

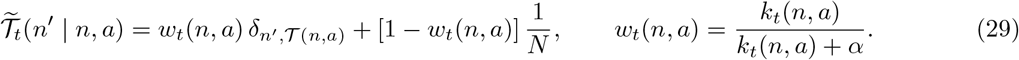

The traversal counts *k*_*t*_ therefore affected planning only through the subjective transition model 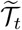; repeated traversals increased the animal’s confidence about where each action led.

#### Planning and policy

At each step, we computed the policy by planning over the *N* = 127 maze nodes using the current rewards and the animal’s estimated transition model:

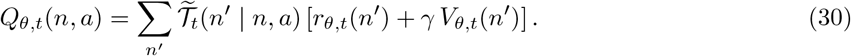

We obtained *V*_*θ,t*_ and *π*_*θ,t*_ using Eqs. 6 and 4, and solved the above equations until convergence for all nodes and actions.

Rewards and transition beliefs were held fixed during planning at each step *t*. They updated after each action, but the planner did not anticipate these future learning updates.

#### Model fitting and selection

For each candidate combination of *λ*_*v*_, *λ*_*u*_, and *α*, we precomputed the experience traces and subjective transition models. We then optimized *θ* with Adam using the conditional action likelihood (Eq. 12), where the policy was computed by dynamic programming over the nodes. We initialized the reward network’s output layer near zero to obtain an approximately flat initial reward.

For each animal, we held out 20% of whole trials within each session and used the same partition for all fits. We evaluated the full hyperparameter grid with five random initializations per configuration. We retained the fit with the highest mean held-out action log-likelihood. The hyperparameter grids, network architecture, and optimization settings are given in Table 3.

#### Analysis of inferred rewards and values

For Fig. 4d, we used the animal’s experience traces and traversal counts at the end of sessions 1, 2, and 10. For each of the two cues shown, we evaluated the reward at every node and solved Eq. 30 for the action values of moves from parent to child nodes. We mean-centered each action-value map over the maze and used a common color scale across sessions.

For Fig. 4g, we evaluated the reward at each of the four ports at every step. For each port, we set the cue to its associated cue and the ITI indicator to zero, regardless of the actual task context. We also computed the mean reward over non-port nodes and all four cues, with the ITI indicator set to zero. We plotted these rewards against cumulative time in the maze on a logarithmic time axis. Shading indicated the s.e.m. across animals.

We also visualized the transition confidence *w*_*t*_(*n, a*) (Eq. 29) over learning in Fig. S9 (averaged across animals in Fig. S9b, for an example animal in Fig. S9c).

#### Model ablations

To determine which components of the reward supported prediction and learning, we fitted three ablated models per animal, each removing one input to the reward network: water history, visitation history, or node depth. We used the hyperparameters selected for that animal’s full model, left the remaining reward inputs, the training partition, and the optimization settings unchanged, and refitted the reward network from scratch.

We evaluated each ablation using held-out LL prediction and generative rollouts. For Fig. 4e, we reported the increase in held-out negative log-likelihood relative to that animal’s full model, so positive values indicate worse prediction after removing the feature. For Fig. 4f, we simulated each ablated model using the same cue sequences, session budgets, and rollout procedure as the full model, with 10 rollouts per model (rollout procedure discussed in detail in Baseline models and evaluation metrics), allowing the histories and transition beliefs it retained to continue updating from its own generated experience. In both panels, circles or curves show individual animals or their mean, and error bars and shading show the s.e.m. across animals.

Finally, to test the contribution of transition learning, we recomputed policies using the true maze transitions without refitting, holding reward parameters and histories fixed (Fig. S9). We reported the increase in held-out NLL obtained with learned rather than true transitions, averaging across seeds within each animal.

### Baseline models and evaluation metrics

#### Baseline models

We compared IRL with models that described behavior through marginal action frequencies, action-transition statistics, or direct mappings from the observed state to the next action without planning. We used the same training and held-out partition for every model. We fitted the simulation and cricket-capture baselines to data pooled across episodes or animals, whereas we fitted the maze baselines separately to each animal, as we did for IRL.

The marginal baseline predicted each action from its overall frequency in the training data, *P* (*a*_*t*_). The Markov model predicted each action from the preceding action, *P* | (*a*_*t*_ *a*_*t−*1_), using transition statistics estimated from the training sequences, similar to how MoSeq [18] models syllable sequences. The hidden Markov model (HMM) allowed action-transition probabilities to vary with a persistent latent behavioral state, *P ∈* (*a*_*t*_ | *a*_*t−*1_, *z*_*t*_), with *z*_*t*_ *∈ {*1, …, *K}*. Following shMoSeq [41], the model learned a baseline action-transition matrix shared across latent states and a state-specific bias for each possible next action. Adding these biases to the baseline logits and applying a softmax yielded *P* (*a*_*t*_ | *a*_*t−*1_, *z*_*t*_).

We evaluated the HMM with *K ∈ {*1, …, 10} latent states in all three settings. We selected *K* = 5 for maze learning and *K* = 10 for cricket capture and simulation. A stickiness parameter *κ* added prior mass to self-transitions, with larger values favoring more persistent latent states. For every combination of *K* and *κ ∈ {*10, 100, 1000}, we ran 100 Gibbs sweeps from each of five random initializations, using a symmetric Dirichlet prior with concentration 1 on the latent-state transition matrix and adding prior mass *κ* to its self-transitions. For each *K*, we selected *κ* using held-out likelihood and retained all five random initializations at the selected value. Fig. S4 shows the full dependence of HMM performance on *K* and *κ*.

We additionally fitted behavior-cloning models that predicted the next action directly from the observed state, *P* (*a*_*t*_ | *s*_*t*_), without representing reward or planning.

For the simulation, BC received the same state representation as IRL. We parameterized it as a multi-layer perceptron with two hidden layers of 128 ReLU units and one output per movement direction (same architecture as that of the Q-network in IRL), with a softmax at the end. We maximized the likelihood of the observed actions using Adam with a learning rate of 3 *×* 10^*−*3^. We fitted five random initializations on the same training trajectories.

For cricket capture, BC again received the same observed state as IRL (Eq. 18), including the preceding syllable, and returned a distribution over the 30 possible next syllables. We parameterized this distribution with a multilayer perceptron (two hidden layers with 128 and 64 ReLU units, with a softmax at the end; same architecture as that of the Q-network in IRL) and trained it by maximizing the action likelihood on the same training trials with a learning rate of 1 *×* 10^*−*3^, fitting five random initializations.

For maze learning, BC took the form of a node–cue lookup table, *P* (*a* | *n, c*), containing a separate action distribution for each maze node and active cue. This model captured location- and cue-dependent choices but did not depend on the animal’s accumulated experience and therefore could not change across sessions. We estimated the table from empirical counts with one pseudocount per available action. Node–cue pairs absent from training received a uniform distribution over available actions and accounted for 0.2% of held-out steps. During inter-trial intervals, we assigned the cue a fifth “no cue” value and scored these steps like all others.

#### Held-out prediction

We held out 25% of complete episodes in the simulation and 20% of complete trials in the cricket-capture and maze datasets, sampling within each session where applicable. We measured predictive performance as the mean negative log-likelihood per held-out action,

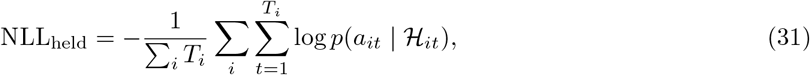

where ℋ_*it*_ contains the current observed state and the animal’s observed history before that action, as in the maze task. For HMM and latent-goal IRL, we computed these probabilities by forward filtering, without using future actions:

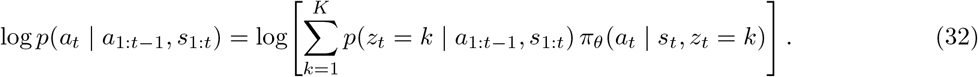

Maze IRL required one additional choice because its policy depended on the animal’s accumulated experience. We masked actions in held-out trials from the fitting objective but continued to update the visitation and water-received traces and transition counts from the animal’s observed experience during those trials. The model’s internal state at each held-out step therefore reflected everything the animal had encountered up to that point. Scoring remained causal without treating held-out trials as experiences the animal had never undergone.

For Figs. 2b and 3b, we fitted IRL, HMM, and BC from five random initializations, repeating the IRL and HMM fits at each value of *K*. Points show individual initializations, bars show their mean negative log-likelihood, and error bars show the s.e.m. across initializations. The marginal and Markov baselines were estimated directly from training counts and did not depend on random initialization, so we show them without error bars across initializations. In Fig. 4b, circles show individual animals, bars show the mean across animals, and error bars show the s.e.m. across animals. We also show individual-animal learning curves in Fig. S8, by averaging IRL rollout performance across five training seeds, with shading indicating the s.e.m. across seeds.

#### Generative behavioral rollouts

We additionally compared the models by simulating them forward in the corresponding environment. We held fitted model parameters fixed during evaluation and used the same environmental dynamics for every model, so that the models differed only in how they selected actions. At each simulation step, we restricted action selection to actions available in the current state. For the marginal baseline, we sampled actions independently from their overall frequency. For the Markov model, we conditioned on the preceding action. For the HMM, we sampled both the evolving latent state and the next action from the fitted model. For BC, we sampled directly from its state-conditioned action distribution. For IRL in simulation and cricket capture, we sampled the initial goal from *p*_0_, evolved it according to the fitted goal-transition model, and sampled actions from the policy associated with the current goal.

For cricket capture (Fig. 3c), we generated *N* = 100 rollouts per model for each of five random seeds for IRL, BC, and HMM. Rollouts began from mouse and cricket configurations matched to held-out trials, using the same initial configurations for every model. Each rollout continued for at most *H* = 300 steps, where one step corresponded to one syllable rather than one video frame. After each sampled syllable, we updated the mouse’s position and heading using the syllable-conditioned movement distributions. All models were therefore evaluated under the same fitted mouse and cricket dynamics. After capture, the rollout entered the post-capture condition and continued until the mouse reached home or exhausted the simulation horizon. Capture occurred when the mouse came within 0.1 normalized arena units of the cricket. Home arrival used the same threshold for the distance between the mouse and home.

We defined a trial as complete only when the mouse had captured the cricket and subsequently reached home. Letting *T*_*i*_ denote the first step at which rollout *i* completed this sequence, we reported

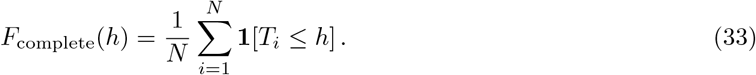

For rollouts that did not complete the trial within *H* steps, we set *T*_*i*_ = *∞*, so they remained in the denominator at every *h*. We computed the empirical curve identically from held-out trials. Curves show the mean across seeds, with shading indicating the s.e.m. across seeds.

For maze learning (Fig. 4c), we generated 10 rollouts for each animal and model. Each rollout spanned all ten training sessions, using the animal’s observed cue sequence and session boundaries. Each simulated session ended after the observed number of node transitions or 250 completed trials, whichever came first. A trial was completed when the model entered the port associated with the active cue. Following each completed trial, the model entered an inter-trial interval lasting the same number of node transitions as the corresponding observed interval. It then advanced to the next cue in the animal’s recorded sequence. No rollout exhausted the recorded cue sequence.

Here, IRL updated its experience traces and transition beliefs after every generated action, allowing its policy to change through simulated experience. The tabular BC model used fixed action probabilities for each node and cue. The Markov and HMM models generated actions using fixed transition parameters, with the HMM additionally sampling transitions between latent states.

For each session, we averaged the number of completed trials first across the 10 rollouts for each animal and then across animals. We report the s.e.m. across animal-level means. Learning curves for individual animals are shown in Fig. S8.

