## Supplemental Figures and Video captions for "Inferring the evolving objectives that organize animal behavior"

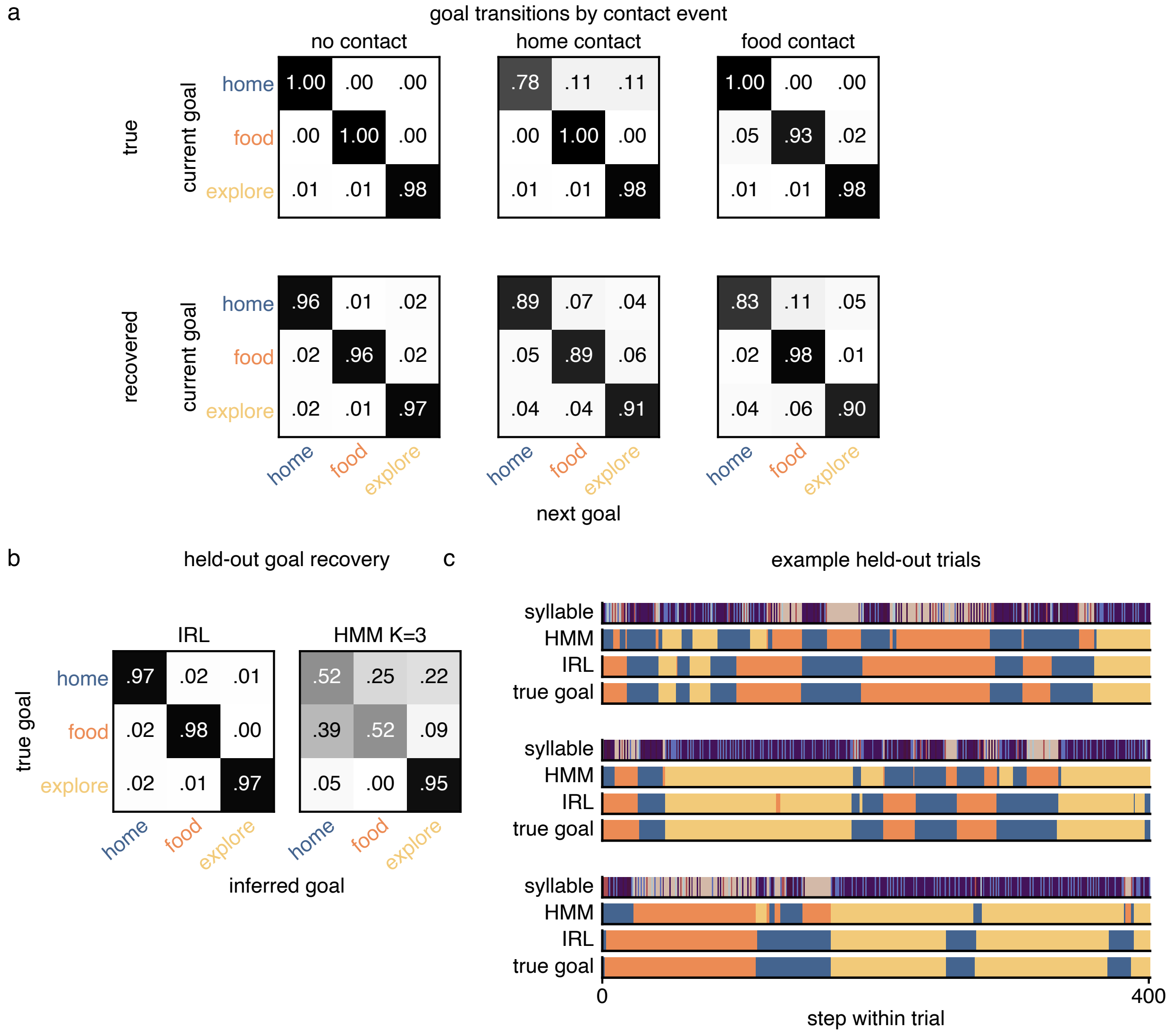


**Figure S1: Recovery of goal transitions and goal sequences in simulation. Related to Fig. 2.** The model was fitted to 150 training trials and evaluated on 50 held-out trials. Inferred states were matched to true goals using the permutation that maximized the mean correlation between inferred and true rewards on held-out states. **(A)** True (top) and inferred (bottom) goal-transition probabilities, conditioned on no contact, home contact, or food contact. Rows indicate the current goal and columns the next goal. **(B)** Row-normalized goal-confusion matrices for IRL and the action-only HMM baseline ($K = 3$). IRL recovers all three goals with 97–98% recall. The HMM identifies exploration from action statistics alone but confuses the two site-seeking goals. **(C)** Actions, inferred states, and true goals for three randomly selected held-out trials.

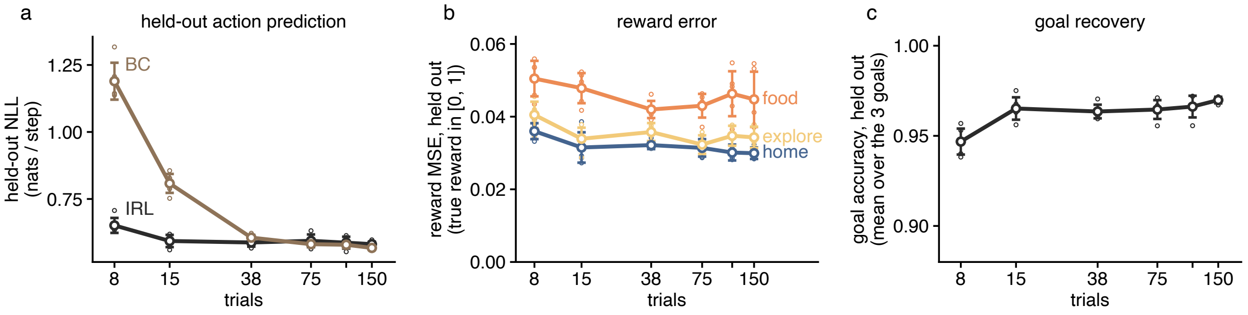


**Figure S2: Data requirements for recovery in simulation. Related to Fig. 2.** Models were fitted to nested subsets of 8–150 trials (400 steps each) and evaluated on the same 50 held-out trials. Model architectures were held fixed across dataset sizes and matched those used in Fig. 2. Open circles show five seeds per subset size; larger markers and bars show the mean ±1 s.d. Trial counts are plotted logarithmically. **(A)** Held-out action negative log-likelihood (NLL). IRL performs well with only eight trials (3,200 time steps), whereas behavior cloning (BC) requires more data to achieve comparable predictive accuracy. **(B)** Mean squared error (MSE) between true and inferred rewards at held-out arrival states. **(C)** Goal accuracy averaged across the three goals, remaining above 94% even with eight trials.

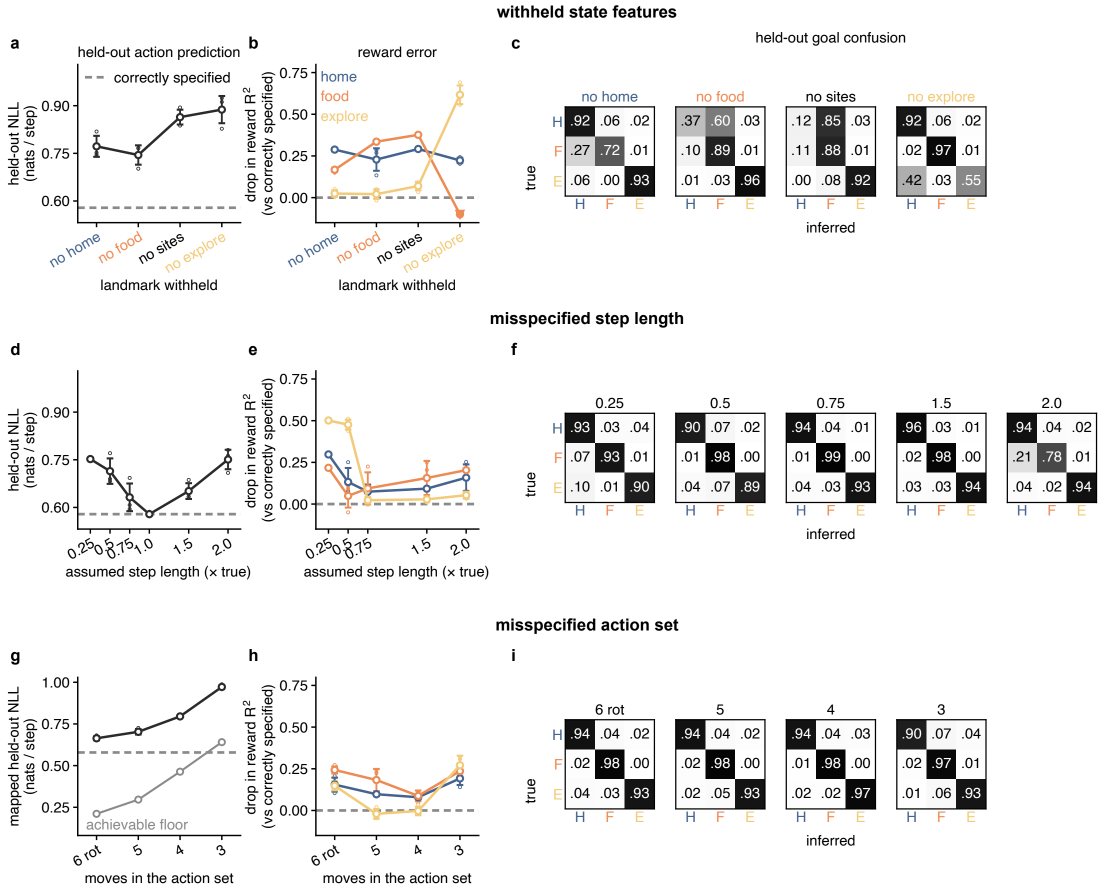


**Figure S3: Effects of model misspecification in simulation. Related to Fig. 2.** We tested reward and goal recovery under several model misspecifications. All conditions used the same 150 training and 50 held-out trials, with three seeds per condition. Each row shows action NLL, reward error (the drop in $R^2$ from the correctly specified fit), and row-normalized goal-confusion matrices (rows, true goal; columns, inferred goal; H, home; F, food; E, explore). Dashed lines mark the correctly specified reference; negative reward error indicates improved recovery. **(A–C)** To test misspecification of the state representation, we removed distance and bearing features for one or both sites, or for the wall, together with allocentric position. Removing site features increases home–food confusion; removing features for both sites largely merges these goals. Removing wall features mainly impairs exploration. These features are therefore crucial for inferring accurate goals and rewards. **(D–F)** To test misspecification of the transition model used for planning, we varied the fitted model's assumed step length from 0.25 to 2 times the true step length of 5.2 cm. NLL is lowest at the true length. Very short assumed steps impair recovery of the exploration reward, but overall goal recovery remains largely intact. Doubling step length produces the largest loss of goal recovery in this test, increasing food-to-home confusion and reducing food-goal recall to 78%. **(G–I)** To test misspecification of the action set, we rotated the model's six moves by 20° or used five, four, or three equally spaced moves, with corresponding forward dynamics. The held-out data still contain the generating model's original six actions. To score these observations, we mapped each action to its nearest model move and divided that move's probability equally among the original actions assigned to it (*mapped NLL*). Fewer moves force distinct original actions to share probability, raising NLL even for the true-policy reference (gray). Despite these action-set misspecifications, recall remains at least 90% for every goal, and the inferred rewards are also close to those of the correctly specified model.

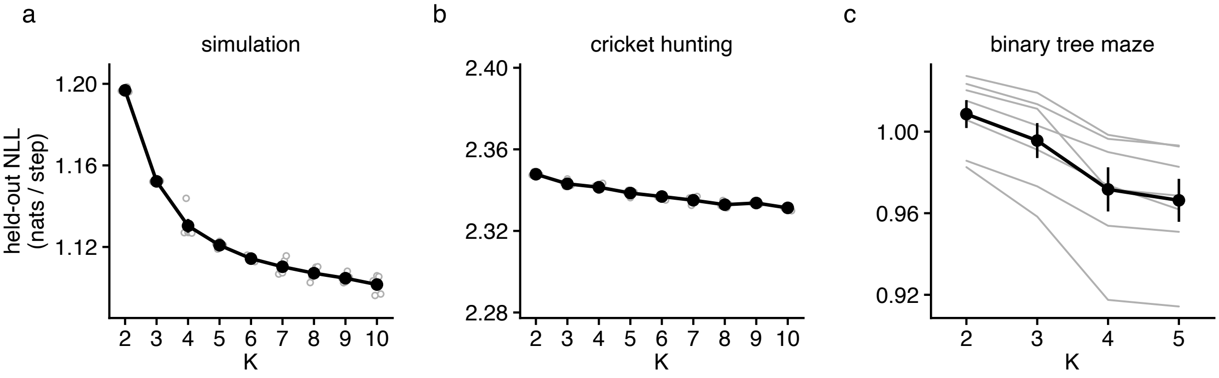


**Figure S4: HMM prediction with different numbers of hidden states. Related to Figs. 2–4.** One-step-ahead held-out action NLL for the HMM baseline with $K$ hidden states and a separate action-transition matrix for each state. For each $K$, the self-transition prior $\kappa \in \{10, 100, 1000\}$ was selected by mean held-out NLL across seeds. **(A)** Simulation. **(B)** Cricket hunting. Open circles show five seeds; larger markers and bars show the mean $\pm 1$ s.e.m. **(C)** Binary tree maze, seven saline control mice. Gray lines show individual mice averaged over five seeds; black markers and bars show the mean $\pm 1$ s.e.m. across mice. Prediction improves with additional states, with diminishing gains; improvements are particularly small in cricket hunting.

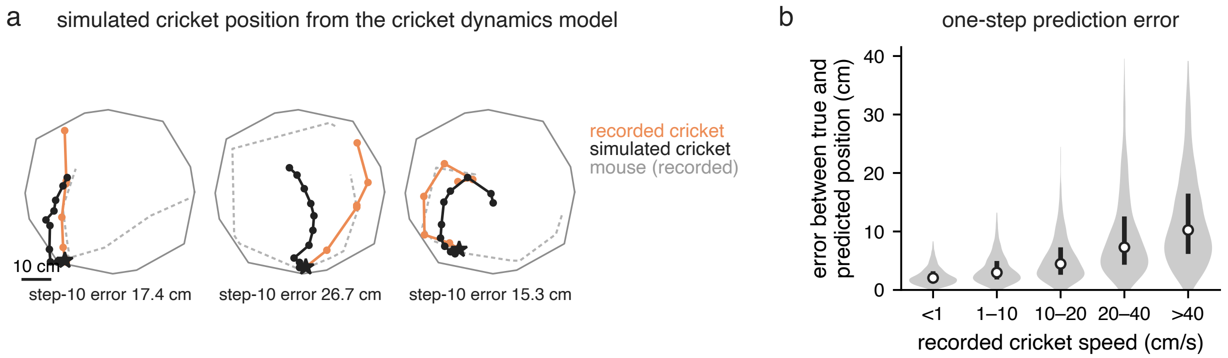


**Figure S5: Validation of the cricket dynamics model. Related to Fig. 3.** A feed-forward network predicts cricket position at the next syllable onset from the current cricket position, mouse position and heading, and syllable. Evaluation uses held-out trial segments before capture. **(A)** Three 10-syllable rollouts with predicted cricket positions fed back into the model and recorded mouse states and syllables supplied as inputs. Recorded cricket trajectories are orange, predictions black, and mouse trajectories dashed gray; stars mark starting positions. Final position errors are shown. **(B)** One-step position error grouped by recorded cricket speed. Violins are truncated at the 99th percentile; white markers and bars show medians and interquartile ranges. Error increases with speed, indicating reduced accuracy for rapid cricket movements.

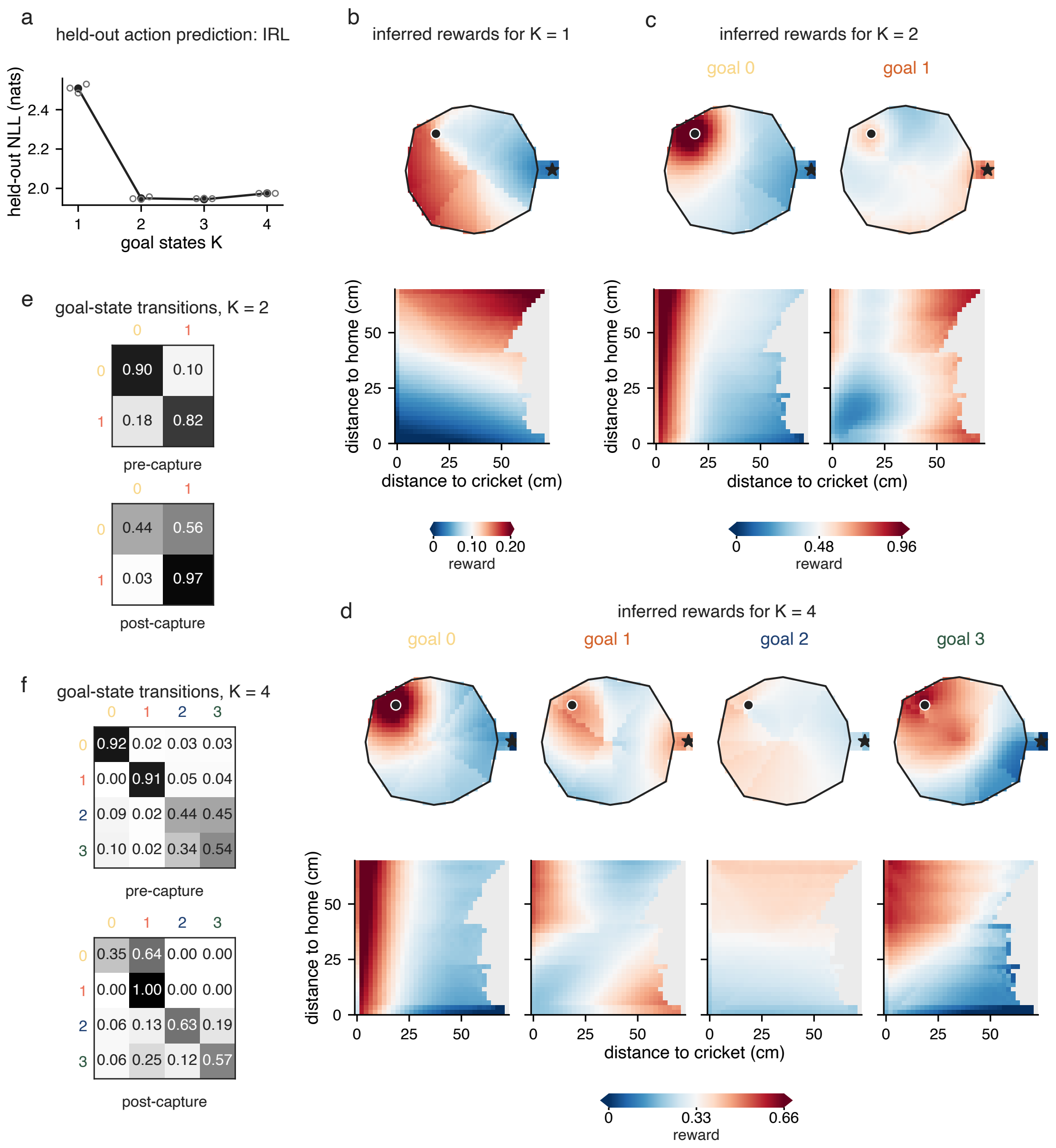


**Figure S6: Number of goal states in cricket hunting. Related to Fig. 3.** IRL models were fitted with different numbers of goal states, $K$, using five seeds per condition. Other hyperparameters were selected by held-out NLL; reward maps and transition matrices are shown for the best fit at each $K$. **(A)** Held-out syllable NLL; open circles show seeds and the line joins their means. Adding a second goal substantially improves prediction, with little further gain at $K = 3$ or 4. **(B–D)** Inferred rewards for $K = 1$, 2, and 4, with one column per goal. Top: arena coordinates with the cricket fixed at the filled circle and home at the star. Bottom: distance to cricket versus distance to home. Rewards are maximized over the next syllable and averaged over heading and previous syllable. Blank regions are geometrically impossible, and color limits are shared across goals within each $K$. At $K = 2$, rewards distinguish pursuing the cricket and movement away from the cricket; $K = 4$ includes a pursuit state that is concentrated on the cricket, a home state that is rewarding at both cricket and home, and two diffuse search states with different reward profiles. **(E, F)** Goal-transition matrices before and after capture for $K = 2$ and $K = 4$ (rows, current goal; columns, next goal). Before capture, the model switches frequently between the two search states at $K = 4$. Capture increases persistence of the home-directed state, whose self-transition probability reaches 0.97 and 1.00 for $K = 2$ and 4, respectively.

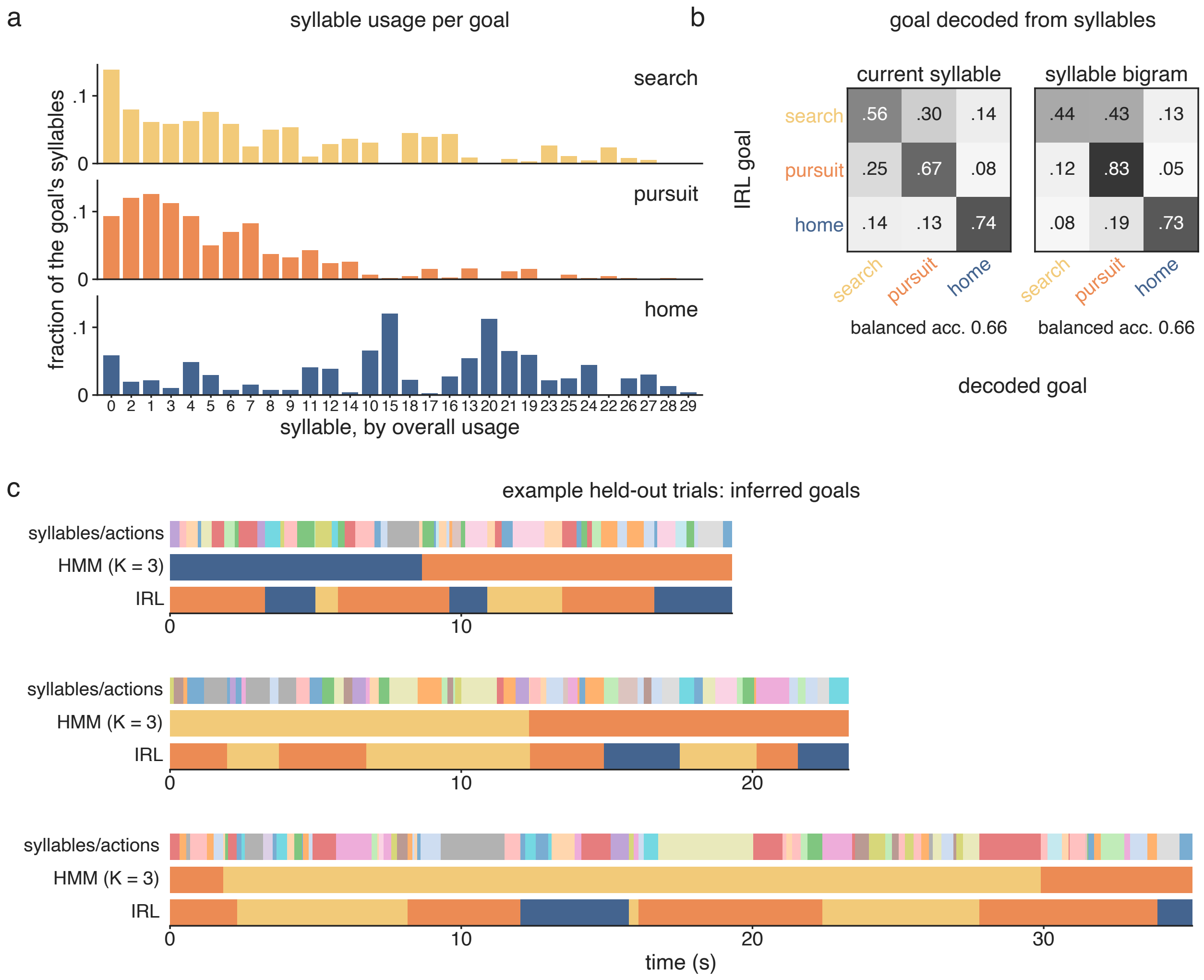


**Figure S7: Relationship between inferred goals and behavioral syllables. Related to Fig. 3.** All panels use the fitted three-goal IRL model from Fig. 3, evaluated on held-out cricket-hunting trials. **(A)** Syllable usage within each goal, weighted by goal posterior probabilities and ordered by overall usage. Search and pursuit have overlapping syllable distributions, while home uses a different mixture of syllables. **(B)** Decoding the most probable IRL goal using multinomial logistic regression trained on either the current syllable alone or the current and previous syllables. Decoders were fitted on training trials; matrices show row-normalized held-out predictions (rows, IRL goal; columns, decoded goal). Both reach 66% balanced accuracy, indicating that local syllable identity only partly predicts the inferred goals. **(C)** Syllables, HMM states ($K = 3$), and IRL goals in three held-out trials selected at the quartiles of HMM–IRL agreement. HMM states were matched to IRL goals on training trials. HMM states persist for longer intervals than IRL goals in these examples.

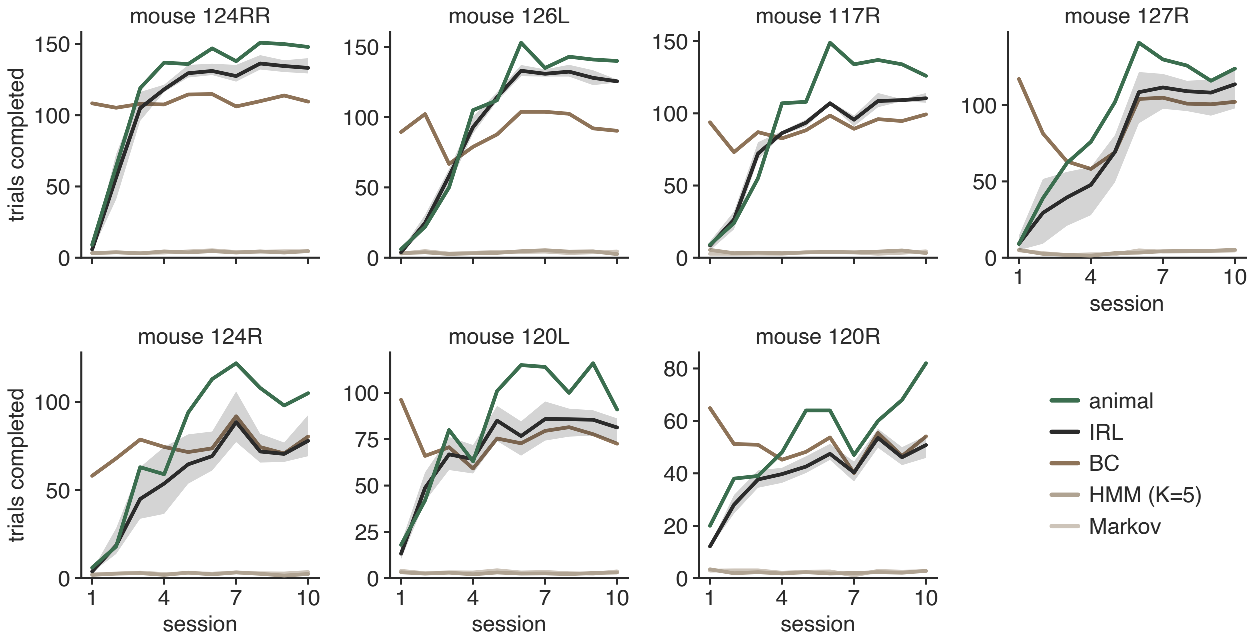


**Figure S8: Maze learning curves for individual animals. Related to Fig. 4.** Trials completed per session by seven mice and in free rollouts of fitted IRL, BC, HMM ($K = 5$), and Markov models. Simulations use each mouse's per-session step budget, and curves show averages across simulated agents. IRL curves additionally average across five training seeds, with shading showing the range across seeds. Mice are ordered by trials completed over the final three sessions; vertical scales vary across panels. IRL captures the increase in performance across sessions. BC overestimates early performance, while the movement-sequence baselines complete few trials.

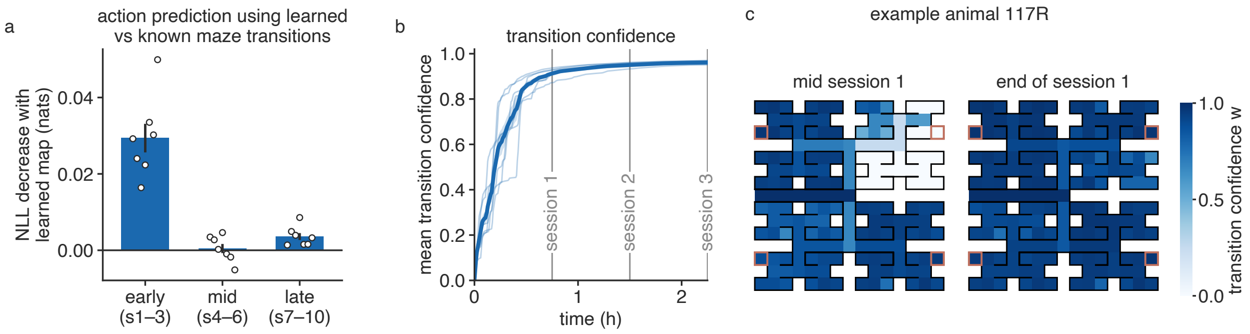


**Figure S9: Learning maze transitions improves early action prediction. Related to Fig. 4.** To model uncertainty about an unfamiliar maze, IRL assigns each edge a transition confidence $w = k/(k + \alpha)$ after $k$ traversals, interpolating between uniform next-state probabilities ($w = 0$) and the true transition ($w = 1$). The resulting transition model is used for planning. **(A)** Increase in held-out NLL when the same fitted model plans with true rather than learned maze transitions, without refitting. Positive values favor learned transitions. Circles show results for seven mice averaged over seeds; bars show the mean $\pm 1$ s.e.m. across mice. The improvement is largest in sessions 1–3 and small thereafter. **(B)** Mean transition confidence versus cumulative time in the maze. Thin lines show mice; the thick line and band show the mean $\pm 1$ s.e.m. Vertical lines mark the ends of sessions 1–3. Confidence approaches saturation by the end of session 2. **(C)** Transition confidence for mouse 117R midway through and at the end of session 1. Confidence increases across much of the maze during the first session. Edge colors indicate confidence; red squares mark reward ports.

### Supplemental video legends

**Video S1. Examples of the inferred search goal.**
**Related to Fig. 3.** A montage of recorded cricket-capture episodes assigned to the search goal by the fitted IRL model, drawing examples from multiple trials and animals.

**Video S2. Examples of the inferred pursuit goal.**
**Related to Fig. 3.** A montage of recorded cricket-capture episodes assigned to the pursuit goal by the fitted IRL model, drawing examples from multiple trials and animals.

**Video S3. Examples of the inferred home goal.**
**Related to Fig. 3.** A montage of recorded cricket-capture episodes assigned to the home goal by the fitted IRL model, drawing examples from multiple trials and animals.

**Video S4. Inferred goal switching during a recorded cricket-capture trial.**
**Related to Fig. 3.** A complete held-out trial with the inferred goal state and behavioral syllables overlaid on the recorded behavior.

**Video S5. A generated cricket-capture trial.**
**Related to Fig. 3.** One trial generated by the fitted switching-goal IRL model in the cricket-capture simulator. The display identifies the simulated mouse, cricket, home location, and active sampled goal as the trial unfolds.

**Video S6. Generated maze trials across learning sessions.**
**Related to Fig. 4.** Generated maze trials from successive learning sessions are concatenated into a single video. The fitted reward parameters remain fixed across sessions, while the model's visitation and water histories and transition beliefs evolve through its simulated experience.
